# Molecular factors shaping small RNA content in mouse sperm

**DOI:** 10.64898/2026.07.27.740910

**Authors:** Giulia Perillo, George Edward Allen, Salman Shehzada, Keigo Shibata, Beatrix M. Ueberheide, Stéphanie Conzelmann-Prin, Marcelle Vanora Darques, Puneet Sharma, Pei-Hsuan Wu

## Abstract

In mammalian reproduction, sperm and oocytes fuse to generate embryos. In contrast to maternally inherited small RNAs, sperm-borne small RNAs have been implicated in embryo viability but remain incompletely understood. Unanswered questions about molecular forces shaping small RNA content in sperm and sources of reporting discrepancies hinder a deeper understanding of their post-fertilization roles. Here, we investigate the dynamics and protein association of small RNAs in C57BL/6 mice during spermiogenesis and sperm capacitation. Using unique molecular identifiers and spike-ins to minimize technical biases in low-input sequencing, we demonstrate a selective sperm small RNA repertoire derived from a subset of spermatogenic small RNAs that remains malleable during capacitation. Retained PIWI-interacting RNAs persist despite widespread RNA decay and are insensitive to capacitation, which may be partly attributed to their association with MIWI proteins. Our data reveal RNA species susceptible to PCR duplicate-related abundance overestimation and uncover biological and technical factors shaping experimentally observed sperm small RNA profiles. These findings refine our understanding of sperm-borne RNAs and inform future functional studies and medically assisted reproduction.

## Introduction

Parental genomes largely dictate the viability and development of a zygote. Beyond genomes, maternally inherited small RNAs, including endogenous siRNAs (endo-siRNAs), microRNAs (miRNAs), and PIWI-interacting RNAs (piRNAs), play a crucial role in development in worms (de Albuquerque *et al*, 2015; Quarato *et al*, 2021; Rieger *et al*, 2023), flies (Barckmann *et al*, 2015; Brennecke *et al*, 2008; Fabry *et al*, 2021; Kugler *et al*, 2013; Rouget *et al*, 2010), and mice (Suh *et al*, 2010; Tang *et al*, 2007). Studies in invertebrates and vertebrates also support a role for sperm-borne small RNAs in offspring development. Life history and environmental factors have been linked to alterations in sperm RNA content and offspring development (Chen *et al*, 2016; Ord *et al*, 2020; Rando, 2012; Rodgers *et al*, 2015; Sharma *et al*, 2016; Wang *et al*, 2021b). tRNA-derived fragments (tRFs) (Chen *et al*, 2016; Sharma *et al*, 2016) and miRNAs (Rodgers *et al*, 2015; Wang *et al*, 2021b) have been proposed as carriers of environmentally induced paternal effects. More broadly, paternal small RNAs have been implicated in embryo development in worms, mice, bulls, and humans as part of the intrinsic program (Alves *et al*, 2019; Boerke *et al*, 2007; Choi *et al*, 2021; Conine *et al*, 2018; Isacson *et al*, 2025; Liu *et al*, 2013, 2012; Rassoulzadegan *et al*, 2006; Stoeckius *et al*, 2014; Wang *et al*, 2021a; Wu *et al*, 2020; Yuan *et al*, 2015). To identify responsible small RNAs and understand their mechanisms, numerous published studies have described the small RNA profiles of mature mouse and human sperm (Chen *et al*, 2016; Hutcheon *et al*, 2017; Krawetz *et al*, 2011; Pantano *et al*, 2015; Peng *et al*, 2012; Sharma *et al*, 2016, 2018; Stanger *et al*, 2020). However, how sperm’s small RNA content is shaped for its post-fertilization role, responds to environmental cues, and transfers information to zygotes remains an open question. An intergenerational function of sperm-borne small RNAs suggests the existence of selective mechanisms, yet their identities are poorly understood. Due to diverse class- and species-specific properties, biogenesis, and turnover pathways of respective small RNA classes (Bartel, 2018; Ghildiyal & Zamore, 2009; Ozata *et al*, 2019), a deeper understanding of their origins and dynamics in sperm may help resolve fundamental questions about paternally inherited RNAs.

In mammalian spermiogenesis, haploid spermatids undergo morphological and molecular transformation characterized by chromatin compaction, cytoplasmic elimination, and progressive shutdown of the transcription and translation machinery (de Kretser *et al*, 1998; O’Donnell *et al*, 2011). Consequently, sperm carry an extremely dense genome and a minute cytoplasmic volume relative to somatic cells, lacking operating gene expression pathways or RNA biogenesis, processing, and base-modification mechanisms. These cells rely on other mechanisms, including signaling pathways and posttranslational modifications of existing proteins, to acquire motility and fertilizing capability during epididymal maturation and to achieve capacitation (Gervasi & Visconti, 2017; Puga Molina *et al*, 2018), both of which are essential for fertilization *in vivo*. The transcriptional quiescence coincides with extensive ribosomal RNA (rRNA) degradation, reflected by the absence of intact 18S and 28S rRNAs that dominate somatic cells (Bianchi *et al*, 2018; Georgiadis *et al*, 2015; Johnson *et al*, 2011). The extent and timing of RNA decay are unclear, which confounds the interpretation of RNA sequencing data from sperm.

Mouse spermatogenesis produces the initial small RNA repertoire in testicular germ cells. The most abundant classes, piRNAs and miRNAs, guide PIWI- and AGO-clade Argonaute proteins respectively to repress expression of spermatogenic mRNAs or transposable elements (Goh *et al*, 2015, 2015; Ozata *et al*, 2019; Reuter *et al*, 2011; Song *et al*, 2014; Walker, 2022; Wu *et al*, 2020; Yuan *et al*, 2015, 2019), thereby ensuring reproductive success. Subclasses of piRNAs—pre-pachytene, hybrid, pachytene piRNAs—are defined by their developmental timing of expression and interact with dedicated PIWI proteins to execute their functions (Aravin *et al*, 2007, 2008; Girard *et al*, 2006; Kuramochi-Miyagawa *et al*, 2008; Li *et al*, 2013). Spermatocytes (4C, 2N) and spermatids (1C, 1N) produce an extraordinary amount of pachytene piRNAs. Their gene regulatory function, which requires their mouse PIWI protein partners, MIWI (PIWIL1) and MILI (PIWIL2), has been well established (Cecchini *et al*, 2026; Choi *et al*, 2021; Deng & Lin, 2002; Li *et al*, 2013; Reuter *et al*, 2011; Wu *et al*, 2020). In rodents, they are enriched in and required only for the male germline due to the missing female-specific *Piwil3* existing in other mammalian genomes (Gong *et al*, 2025, 2026; Hasuwa *et al*, 2021; Ishino *et al*, 2021; Loubalova *et al*, 2021; Lv *et al*, 2023; Roovers *et al*, 2015; Zhang *et al*, 2021).

In addition to spermatogenesis, two windows of opportunity enable further alterations of sperm small RNA content. The first, epididymal maturation of sperm, has been demonstrated. After their release from the lumen of testicular seminiferous tubules, sperm cells transit through the epididymis to reach the distal (cauda) section, during which sperm acquire epididymal tRFs and miRNAs *via* epididymosomes that subsequently replace the dominant testicular piRNAs (Sharma *et al*, 2016, 2018). Approximately 15–80% of small RNAs are estimated to derive from tRNAs in mature sperm across studies (Chen *et al*, 2016; Hutcheon *et al*, 2017; Peng *et al*, 2012; Schuster *et al*, 2016; Sellem *et al*, 2020; Sharma *et al*, 2016, 2018; Short *et al*, 2017; Stanger *et al*, 2020; Tomar *et al*, 2024; Wang *et al*, 2023). The second, less investigated window of opportunity is sperm capacitation. *In vivo*, capacitation entails further maturation of sperm in the female reproductive tract, which renders hyperactivated motility and global protein tyrosine phosphorylation, among other hallmarks that potentiate sperm’s ability to fertilize (Puga Molina *et al*, 2018; Visconti *et al*, 1998). Despite the physiological necessity of capacitation for fertilization, studies of the capacitated sperm RNA repertoire remain scarce. Data suggest that capacitation further modifies the sperm small RNA landscape in boars (Li *et al*, 2018) and possibly other mammals. However, this hypothesis has not been formally tested.

Here, we report the dynamics and protein association of small RNAs in mature sperm from the common laboratory mouse strain C57BL/6 during spermiogenesis and sperm capacitation. Amidst widespread and continued RNA decay, subsets of intact spermatogenic small RNAs are selectively retained in sperm. Capacitation evokes further mRNA degradation but paradoxically increases the abundance of specific miRNAs and tRFs. Retained spermatogenic pachytene piRNAs remain consistent even throughout capacitation, which may be partly explained by their continued association with MIWI proteins. Incorporating unique molecular identifiers (UMIs) and spike-in oligos in our sequencing analyses uncovers small RNA species prone to artifactually inflated quantification due to PCR duplicates, helping reconcile reporting discrepancies. These observations elucidate biological and technical factors shaping the experimentally observed sperm RNA repertoire and help refine interpretations of paternal contribution studies and outcomes of medically assisted reproductive procedures.

## Results

### Small RNAs are selectively retained in mature mouse sperm

We sequenced 18–36 nt RNAs in caudal epididymal sperm (henceforth, cauda sperm) from C57BL/6 mice using an established procedure incorporating UMIs and spike-in oligonucleotides (Fu *et al*, 2018; Wu *et al*, 2020) (**Table 1** and **Supplementary Table 1**). Sperm were purified using a step gradient to preserve membrane integrity and RNA profiles (Johnson *et al*, 2015; Pearson *et al*, 2018) and chemically and mechanically lysed to maximize RNA yield (**Figs. S1A–S1B**). RNA-seq analyses of purified cells showed a higher abundance of previously reported sperm transcripts (Sun *et al*, 2021) than cauda epididymis-expressed transcripts (Shi *et al*, 2021a), validating sperm purity (**Fig. S1C**).

Compared to spermatids, spike-in-normalized small RNAs in sperm exhibited a pronounced increase in rRNA fragments (2.0±0.7% to 53±5.6%; **Fig. 1A**), which likely reflected extensive rRNA decay (Bianchi *et al*, 2018; Georgiadis *et al*, 2015; Johnson *et al*, 2011). In contrast, piRNA abundance fell substantially (91±1.2% to 22±4.8%). Consistent with sperm RNA remodeling during epididymal maturation (Sharma *et al*, 2018), subsets of spermatid miRNAs and tRFs, but not piRNAs, increased in sperm (**Fig. 1B**). Retained piRNA composition closely resembled that of spermatids with predominantly pachytene piRNAs (87.6% of all piRNAs). These sequences carried signatures of canonical pachytene piRNAs, including a ∼30 nt length and a bias for uridine at the first position (**Figs. 1C** and **S1D**). Among them, 46.2% were derived from the ten genomic loci that also accounted for half of spermatid pachytene piRNAs (**Fig. 1D**). The most abundant sequences consistently originated from similar genomic locations within individual piRNA-producing loci between spermatids and cauda sperm, as demonstrated by the piRNA mapping profiles across *pi17* (chr17:27,507,249–27,586,457, encompassing 17-qA3.3-27636 (−) and 17-qA3.3-26735 (+)), the most productive pachytene piRNA-producing locus in adult mouse testes (Aravin *et al*, 2007; Gainetdinov *et al*, 2023; Goh *et al*, 2015; Wu *et al*, 2020) (**Fig. S1E**). Together, the data supported that sperm-borne piRNAs are of germline origin, representing a subset of the spermatogenic pool.

**Figure 1.**
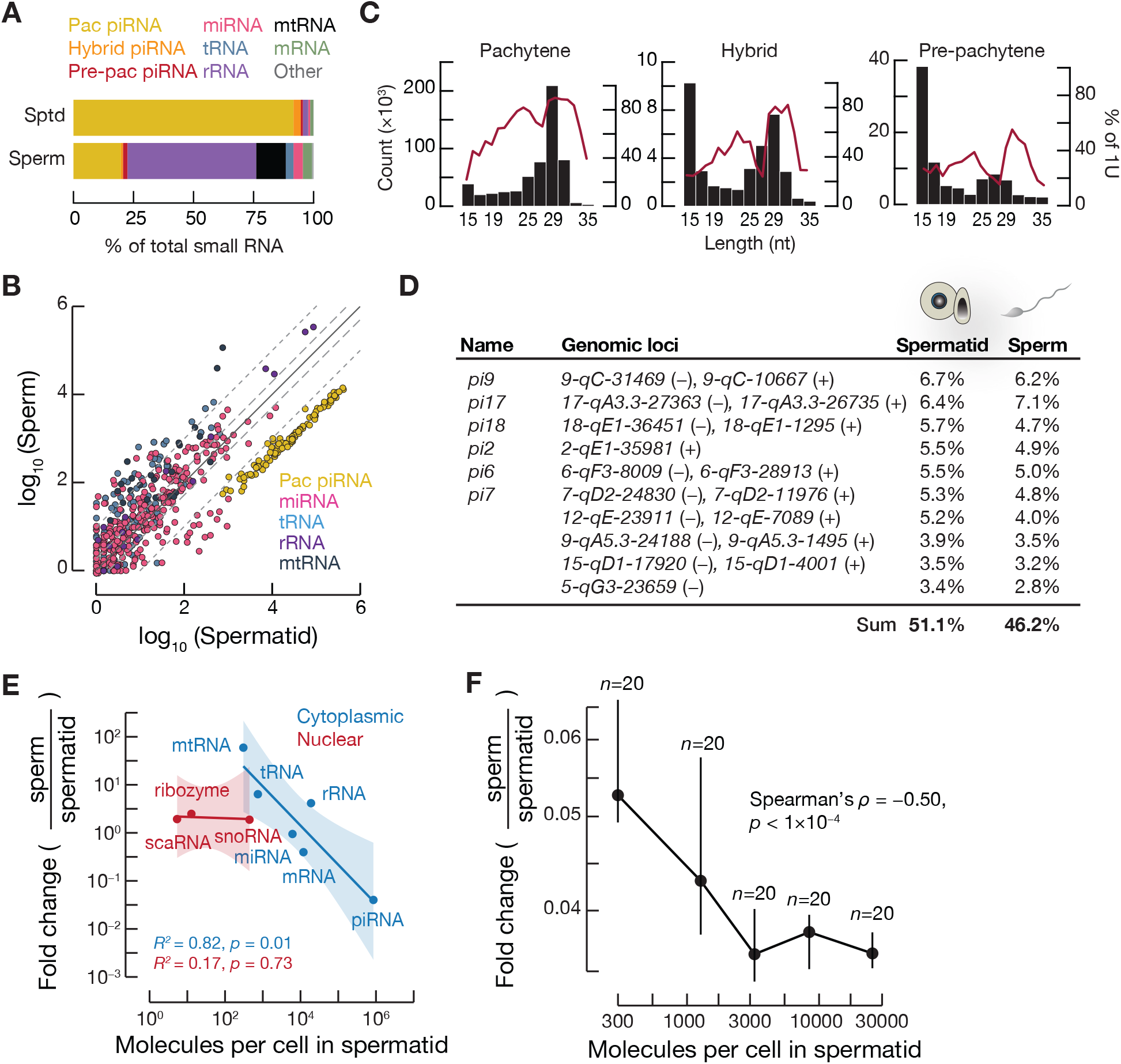
Small RNA composition in mature mouse sperm. **(A)** Small RNA compositions in C57BL/6 spermatids (*n*=3) and caudal epididymal sperm (*n*=3). Sptd, spermatid. Pac piRNA, pachytene PIWI-interacting piRNA. Pre-pac piRNA, pre-pachytene piRNA. miRNA, microRNA. mtRNA, mitochondrial RNA. **(B)** Comparison of small RNAs in C57BL/6 spermatids and sperm. Rare RNA classes are omitted for clarity. Each point denotes the mean spike-in-normalized value averaged across biological replicates with a pseudo-count of 0.5 added before the log transformation (*n*=3 per cell type). Dashed lines mark the 2- and 10-fold changes, respectively. **(C)** Length distribution (histogram) and uridine enrichment at the first position (1U, red line) of piRNA subclasses detected in sperm. **(D)** Fractions of the ten most abundant pachytene piRNA pools in spermatids and their corresponding representation in sperm. Mean piRNA fractions relative to total pachytene piRNAs are shown (*n*=3 per cell type). Locus pairs sharing a bidirectional promoter are presented as a single locus. Published alternative locus names are indicated when applicable. **(E)** Relationship between small RNA abundance in spermatids and retention in sperm. Each point represents the spike-in-normalized value of all non-zero RNAs in the indicated category, averaged across biological replicates in spermatids (*n*=3) and sperm (*n*=3). Shaded regions indicate 95% confidence intervals of the fitted regression lines. Subcellular localization assignments reflect the predominant patterns. **(F)** Relationship between abundance of pachytene piRNA pools in spermatids and their retention in sperm. One hundred defined pachytene piRNA-producing loci are binned by the abundance of piRNAs they produce in spermatids. Each point represents the median values of 20 loci (*n*=3 per cell type). Vertical lines denote the interquartile range.

Similar to pachytene piRNAs, sperm-borne miRNAs harbored the ∼22 nt length profile of intact species, indicating their integrity and authenticity. In comparison, pre-pachytene piRNAs and a subset of hybrid piRNAs could be detected in sperm but at a low abundance. These sequences were markedly shorter than the expected ∼26–30 nt lengths and lacked the characteristic 5′ uridine bias (**Fig. 1C**), suggesting degraded products. We hereafter excluded these categories from downstream analyses.

The abundance of small RNAs in spermatids was inversely correlated with their retention in sperm (**Fig. 1E**). The trend was maintained after removing piRNA or mitochondrial RNA (mtRNA) fragments—which represented the two extremes of the observed abundance-retention relationship—albeit with weaker correlations (**Fig. S1F**). By contrast, nuclear-enriched small RNAs showed no such relationship despite efficient sperm lysis during RNA extraction (**Fig. S1A**), confirming that the cytoplasm is the main site of small RNA remodeling in sperm. piRNAs, which comprised most spermatid small RNAs, also accounted for the greatest RNA decrease in sperm. By examining individual pachytene piRNA pools produced by distinct loci, we observed that most populations also reduced proportionally to their initial abundance in spermatids (**Fig. 1F**). That is, except for the 20 most abundant populations, which deviated from the trend and were preferentially retained in sperm. Together, these results indicate that small RNAs are not uniformly depleted from the cytoplasm at spermiogenesis or randomly degraded at epididymal maturation.

### PCR duplicates artifactually inflate the estimated abundance of tRNA fragments

We noted markedly fewer tRFs in our cauda sperm data (2.6±0.4% of total small RNA) than several reported fractions (∼50–80% of total small RNA) (Hutcheon *et al*, 2017; Schuster *et al*, 2016; Sharma *et al*, 2016; Stanger *et al*, 2020; Tomar *et al*, 2024) despite prioritized tRF assignment over all RNA classes except spike-ins and rRNAs during sequencing read alignment. Because tRNAs are multicopy genes with highly redundant sequences, the observation was at first attributed to the read alignment parameters, which were expected to capture some, but not all, multi-mapped reads (Methods). However, permissive alignment criteria increased estimated tRF abundance only modestly (13±2% of total sperm small RNA; **Fig. S2A**), suggesting that additional factors contributed to the discrepancy between our and published data.

We compared our UMI-incorporated dataset before and after PCR duplicate removal with published small RNA sequencing data from cauda sperm of the same mouse strain but without UMIs (Chen *et al*, 2016). The analysis revealed that PCR duplicates explained much of the difference in estimated tRF abundance. For example, the most abundant tRFs derived from *tRNA-Gly-GCC* and *tRNA-Glu-CTC* in both datasets were comparable when all biologically and artifactually identical sequences were collapsed by non-redundantly counting expressed tRNA loci (**Fig. 2A**; left; **Fig. S2B**). In contrast, UMI-based deduplication, which selectively removed technically identical sequences, reduced *tRNA-Gly-GCC* and *tRNA-Glu-CTC* fragments by approximately tenfold (**Fig. 2A**; middle and right).

**Figure 2.**
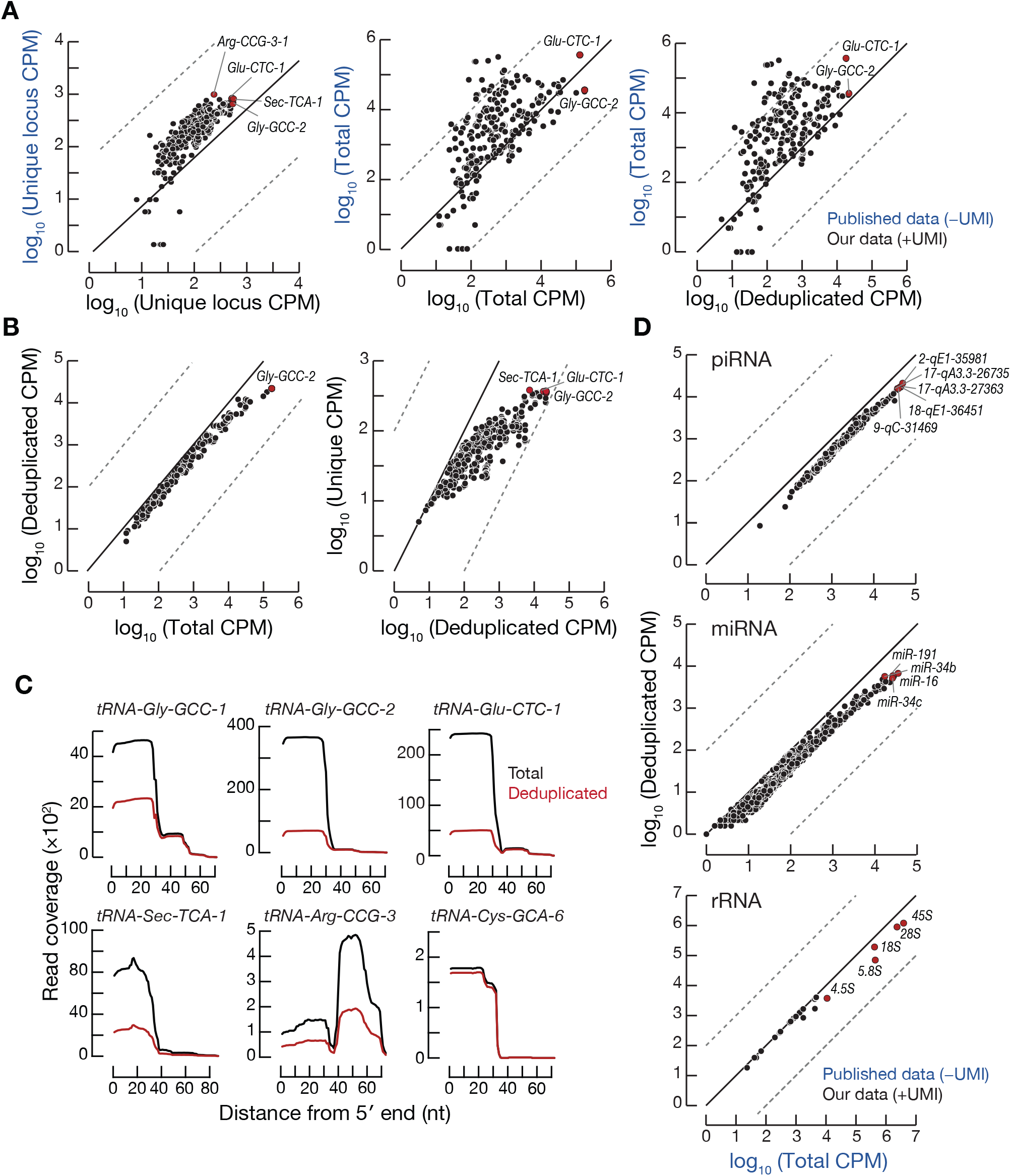
Influence of PCR duplicates on small RNA abundance estimates. **(A)** Estimated tRF abundance in our dataset and published data generated from C57BL/6 cauda sperm. Each dot represents the mean CPM (counts per million) value of fragments derived from a tRNA gene (*n*=3). The most abundant species are highlighted in red. Dash lines indicate 100-fold differences. **(B)** Estimated tRF abundance after removing duplicates using UMIs (left) or collapsing all identical sequences regardless of origin (right). Each dot represents the mean abundance of fragments derived from a tRNA gene (*n*=3). Dashed lines indicate 100-fold differences. **(C)** Read coverage of representative tRNA genes in cauda sperm. Lines represent mean counts across biological replicates (*n*=3). **(D)** Estimated abundance of piRNA, miRNA, and rRNA fragments in cauda sperm from our and a published dataset. Each dot represents the mean abundance of small RNAs produced from a locus (*n*=3). The most abundant species are highlighted in red. Dashed lines denote 100-fold differences.

The most abundant sperm tRFs in both datasets originated from a small subset of loci, including *tRNA-Glu-CTC* and *tRNA-Gly-GCC* (Sharma *et al*, 2016; Wang *et al*, 2023) (**Fig. S2C**). Using UMIs to discern biological molecules from amplification artifacts in our dataset, we found that these highly abundant tRFs were also most strongly impacted by deduplication (**Fig. 2B**). Analysis of individual tRNA loci supported this trend, revealing that tRNAs generating larger numbers of biologically identical fragments from either 5′ (*e.g.*, *tRNA-Gly-GCC*, *tRNA-Glu-CTC*, and *tRNA-Sec-TCA*) or 3′ (*e.g.*, *tRNA-ARG-CGG*) portions of the gene were associated with more PCR duplicates (**Fig. 2C**).

Unlike tRNAs, mouse pachytene piRNAs and miRNAs carry largely non-redundant sequences (Aravin *et al*, 2006; Bartel, 2004; Piriyapongsa *et al*, 2007). Consequently, their quantification was minimally affected by the read alignment parameters (piRNAs: 29±5.1% under restrictive vs 24±3.9% under permissive alignment; miRNAs: 5±1.1% under restrictive vs 5±1.2% under permissive alignment; **Fig. S2A**). In accordance with this, sperm piRNAs, miRNAs, and rRNA fragments (35±4.9% under restrictive vs 27±3.9% under permissive alignment) showed strong agreement between our and the published data regardless of UMI-based deduplication in our analysis (**Fig. 2D**). Nevertheless, similar to tRFs, the most abundant species remained disproportionately affected by PCR duplicates (**Figs. 2D** and **S2D**). For example, *miR-34b/c*, *miR-16*, and *miR-191* displayed some of the largest reductions in estimated abundance following deduplication. Together, these results identify PCR duplicates as a major source of variation in the quantification of sperm small RNA. Highly abundant small RNAs and fragments, particularly those derived from multicopy loci with extensive sequence redundancy, are most susceptible to duplicate-driven inflation.

### Sperm capacitation alters small RNA composition beyond epididymal maturation

To determine whether sperm capacitation influences small RNA composition, we characterized sperm-borne small RNAs before and after *in vitro*-induced capacitation (Boitrelle *et al*, 2021; Quinn *et al*, 1985; Wu *et al*, 2020) (**Fig. 3A**). As expected, incubation under capacitating conditions for 90 min increased protein tyrosine phosphorylation, a hallmark of capacitation (Visconti *et al*, 1995a, 1995b) (**Fig. S3A**). Capacitation elicited no major shift in sperm small RNA composition (**Fig. 3B**) and only moderate (< 2-fold), though statistically significant, changes in four RNA classes across biological replicates with consistent overall small RNA profiles (**Figs. 3C** and **S3B**): pachytene piRNAs (FC = 0.6, *q* < 0.001), tRFs (FC = 1.5, *q* < 0.001), miRNAs (FC = 0.85, *q* = 0.021), and mRNA fragments (FC = 1.2, *q* < 0.001).

**Figure 3.**
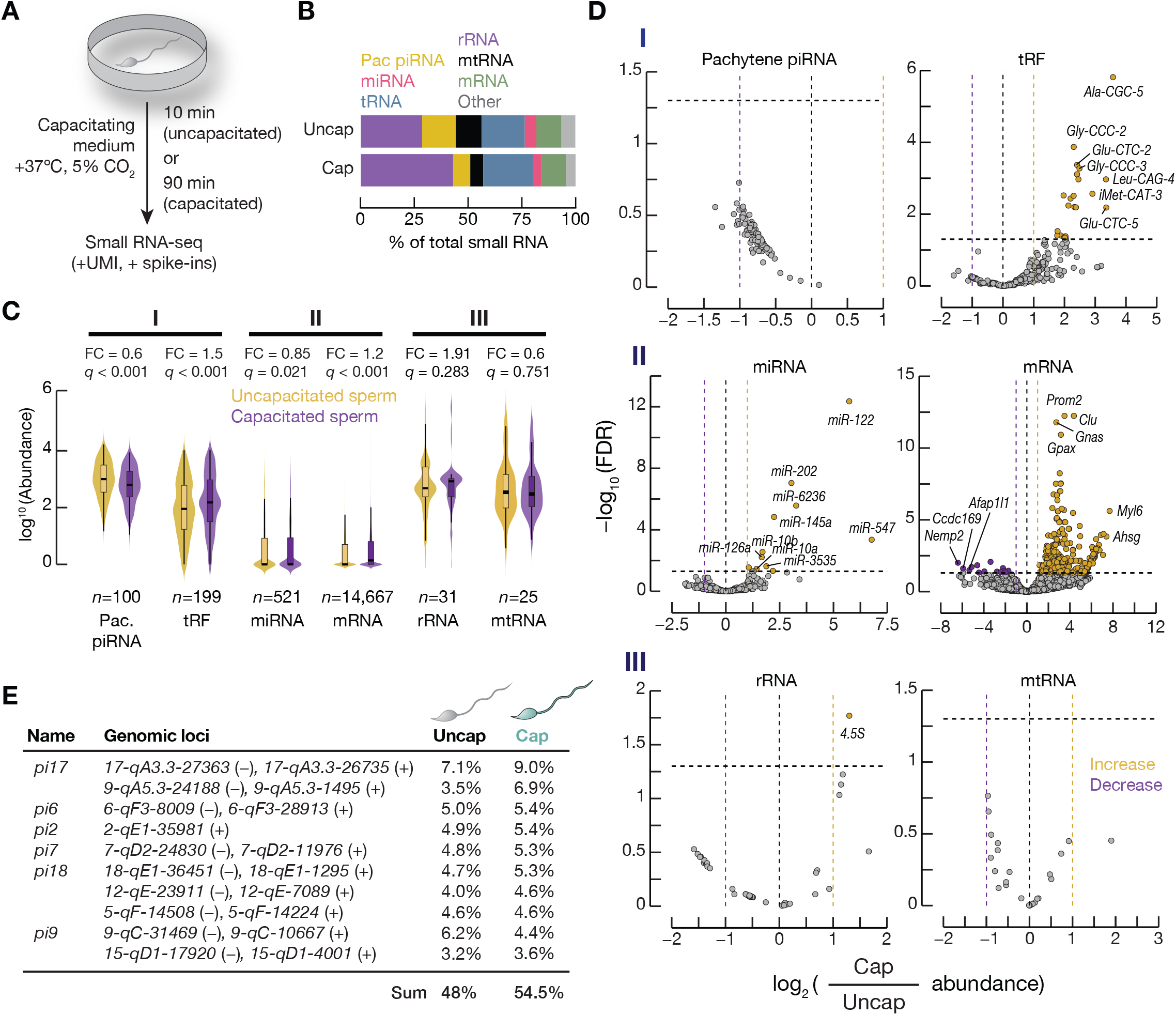
Sperm capacitation influences a subset of small RNAs. **(A)** Schematic of an *in vitro* sperm capacitation assay. **(B)** Relative small RNA abundance in uncapacitated (uncap, *n*=3) and capacitated C57BL/6 cauda sperm (cap, *n*=3). Pac. piRNA, pachytene piRNA. **(C)** Spike-in-normalized mean RNA abundance in uncapacitated (*n*=3) and capacitated sperm (*n*=3) shown as violin plots with boxplot overlays. Boxes indicate the interquartile range, horizontal lines denote medians, and whiskers extend to the maximum and minimum values. RNAs with zero reads across all replicates were excluded. A pseudo-count of 1 was added prior to the log transformation. *q*-values are FDR-adjusted *p*-values from a Wilcoxon rank-sum test. FC, fold change. **(D)** Capacitation-associated changes in specific small RNAs in Category I (top), Category II (middle), and Category III (bottom). Each point denotes the mean spike-in-normalized abundance of RNAs (*n*=3) produced from a distinct locus. **(E)** Fractions of the ten most abundant pachytene piRNA pools in capacitated sperm and their corresponding representation in uncapacitated sperm. Mean fractions relative to total pachytene piRNAs are shown. Locus pairs sharing a bidirectional promoter are presented as a single locus. Published alternative locus names are indicated when applicable.

Despite the overall stable composition, a subset of sperm-borne small RNAs nevertheless responds to capacitation. The six most abundant small RNA classes could be grouped into three categories based on their total abundance and sensitivity to capacitation. Examination of individual RNA species in respective classes revealed those underlying the modest class-level changes induced by capacitation (**Fig. 3D** and **Table 2**). Category I comprises abundant, capacitation-sensitive pachytene piRNAs and tRFs. Increased tRF abundance at the class level was driven by fragments derived from just 5.4% of isodecoders detected in sperm, corresponding to eight isotypes: *Gly*, *Glu*, *His*, *Val*, *Ile*, *Ala*, *Leu*, and initiator *Met*. In contrast, the moderate collective decrease in pachytene piRNAs reflected a concordant but individually statistically nonsignificant reduction across nearly all pachytene piRNA pools. These sequences preserved their characteristic size distribution and 5′-uridine bias following capacitation (**Figs. S3C** and **S3D**), indicating that the remaining sequences were unlikely products of RNA decay. Consistent with this interpretation, pachytene piRNAs identified in uncapacitated and capacitated sperm are highly concordant, with 54.5% of the population in capacitated sperm originating from the same ten loci contributing to 48% of pachytene piRNAs in uncapacitated sperm (**Fig. 3E**).

Category II contained capacitation-sensitive but significantly less abundant miRNAs and mRNA fragments, among which just a subset of sequences were influenced by capacitation (**Figs. 3C** and **3D**, middle panel). Thirteen miRNAs increased after capacitation, representing 11 seed families (*miR-547*, *miR-122*, *miR-6236*, *miR-202*, *miR-145a*, *miR-378*, *miR-3535*, *miR-10a/10b*, *miR-126a*, *miR-143*, *miR-125a*, *miR-143*) and including the *miR-143/145* genomic cluster in the mouse chromosome region 18qE1. Among these, *miR-10a* and *miR-122* have implicated roles in spermatogenesis. Overexpression of *miR-10a* leads to male sterility in mice (Gao *et al*, 2019). *miR-122* is associated with abnormal chromatin biology in sperm differentiated from human induced pluripotent stem cells and with bovine sperm motility (Capra *et al*, 2017; Liu *et al*, 2013). Furthermore, *miR-202* has been proposed to paternally contribute to bovine embryo development (Wang *et al*, 2021a). It is noteworthy that *miR-34b/c*, whose paternal contribution to early embryonic development has been reported but remains inconclusive (Liu *et al*, 2012; Yuan *et al*, 2015), was unaffected by capacitation.

Compared to all examined small RNA classes, mRNA fragments appeared highly responsive to capacitation, accumulating sequences derived from 268 mRNAs. However, the vast majority of mRNA fragments were scarce (**Fig. S3E**). Many increased fragments originated from transcripts encoding ribosomal proteins (GO:0022626, cytosolic ribosome; log_10_(observed/expected) = 1.27, FDR = 5.3×10^−21^; **Fig. S3F**), an observation consistent with extensive rRNA decay in sperm and the notion of prevention of inappropriate translation after spermiogenesis (Johnson *et al*, 2011; Lei *et al*, 2021). Fragments derived from non-ribosomal transcripts were heterogeneous and lacked consistent genomic origins, suggesting no specific processing mechanism. Together with the poorly defined function of mRNA fragments, the observed pattern most likely reflects continued mRNA decay during capacitation.

Category III includes rRNA- and mtRNA-derived fragments (**Figs. 3C** and **3D**, bottom panel). Despite a substantial rise from spermiogenesis to epididymal maturation presumably due to RNA decay (**Fig. 1A**), these small RNAs further accumulated little during capacitation, except fragments from the 3′ end of 4.5S rRNA (*M27443.1*). This suggests that rRNAs and mtRNAs in sperm are degraded mainly before capacitation in the male reproductive tract. Together, these findings demonstrate that capacitation does not substantially remodel the sperm small RNA landscape but selectively increases specific small RNA species, particularly a subset of tRFs and miRNAs. Notably, no specific miRNA, tRF, or pachytene piRNA showed a significant reduction during capacitation, suggesting RNA stability.

### Retained sperm piRNAs are protein-associated

The preferential and consistent retention of small RNAs in uncapacitated and capacitated sperm suggested mechanisms that confer the selectivity or RNA stability to this subset of small RNAs. For example, persisting pachytene piRNAs could reflect their extreme initial abundance in spermatogenesis, stabilization by 3′ terminal 2′-*O*-methylation, continued association with MIWI or MILI proteins, or a combination of these factors. Similarly, continued association of miRNAs with AGO Argonautes beyond spermatogenesis may stabilize miRNAs in sperm (Diederichs & Haber, 2007; Winter & Diederichs, 2011). However, plentiful and well defined in mouse spermatogenic cells, the presence of these proteins in sperm is poorly characterized. In particular, detection of sperm-borne MIWI (Hutcheon *et al*, 2017) is contradicted by the observation that MIWI removal in late spermiogenesis is essential for sperm formation (Gou *et al*, 2017; Li *et al*, 2024; Zhao *et al*, 2013). If MIWI is indeed absent from sperm, stable pachytene piRNAs would be attributed to other explanations. To resolve this possibility, we surveyed piRNA pathway proteins in C57BL/6 uncapacitated sperm by isobaric labeling-based tandem mass tag (TMT) mass spectrometry. C57BL/6 spermatids served as positive controls, in which 14 known piRNA pathway proteins—including MIWI and MILI—and all mouse AGO Argonautes (AGO1–4) (González-González *et al*, 2008; Modzelewski *et al*, 2012) were predictably detected (*n*=5; **Fig. 4A** and **Table 3**). In contrast, MIWI, VASA (DDX4), and TRDR6, but no other AGO, PIWI, or piRNA pathway proteins, were detected in sperm, identifying MIWI—but not MILI—as the tentative protein binding partner of pachytene piRNAs in sperm. Furthermore, this observation predicts that miRNAs do not have a protein binding partner in sperm.

**Figure 4.**
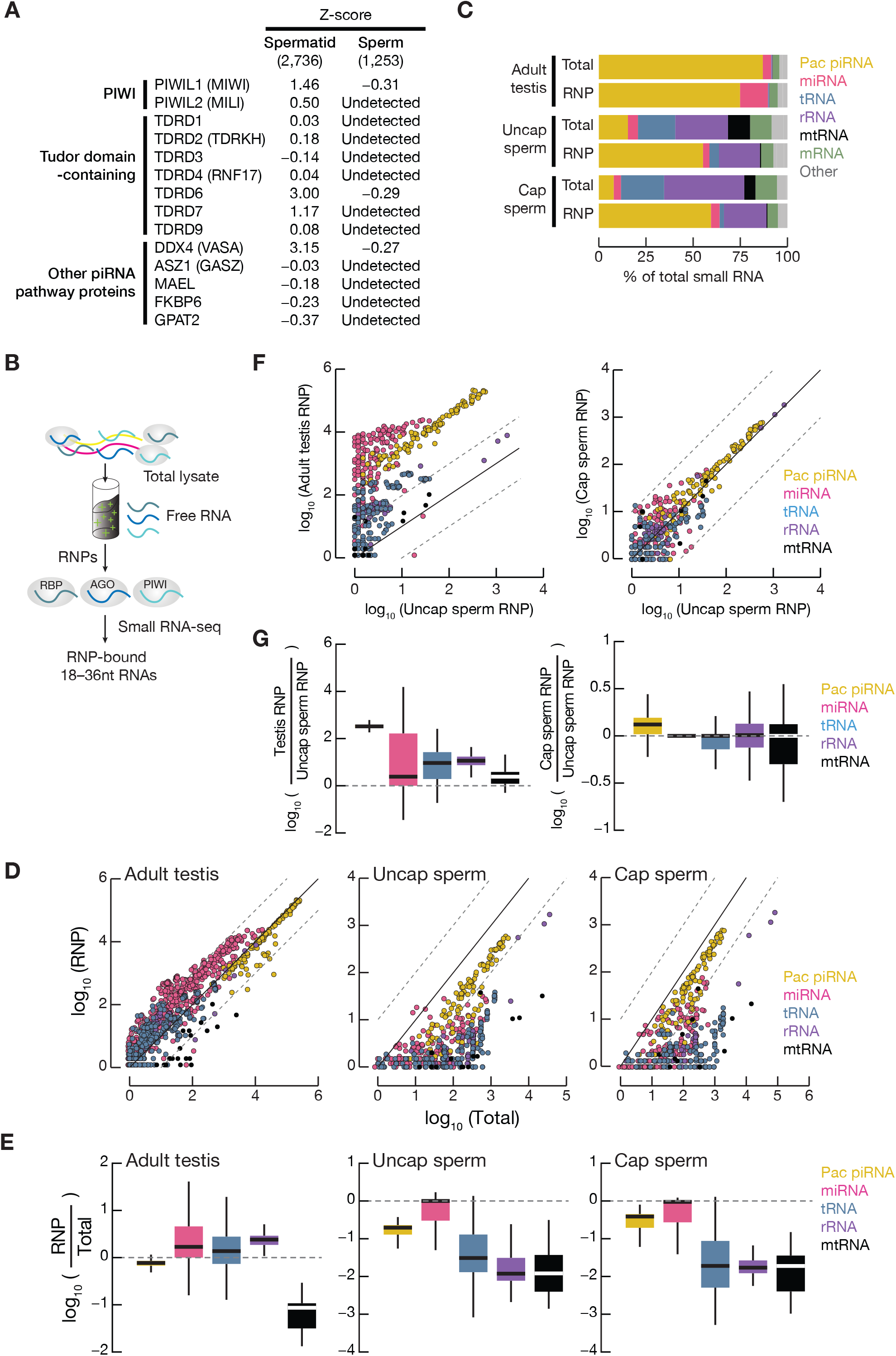
Protein association of small RNAs in mouse sperm. **(A)** piRNA pathway proteins in C57BL/6 spermatids (*n*=5) and uncapacitated cauda sperm (*n*=5). Proteins were identified by tandem mass tag (TMT) mass spectrometry. Respective Z-scores, indicative of the relative protein abundance between spermatids and sperm, are shown (Methods). **(B)** Schematic depicting the ribonucleoprotein (RNP)-bound small RNA isolation procedure using TraPR (Grentzinger *et al*, 2020). **(C)** Relative abundance of RNP-associated small RNAs in adult mouse testes (control, *n*=2) and sperm (*n*=3). RNP-associated small RNAs were purified by TraPR. The “other” category includes snoRNA, lncRNA fragments, ribozyme, and scaRNA. Total, total small RNAs. RNP, RNP-associated small RNAs. **(D)** Comparison of RNP-associated and total small RNAs in adult mouse testes and sperm. Each point represents the mean spike-in-normalized abundance of small RNAs derived from one locus across biological replicates. A pseudo-count of 0.5 was added before log transformation. Spike-in oligos were added after RNA extraction. The solid and dashed lines denote the identity line and ±10-fold fold-change line, respectively. **(E)** Ratios of the mean abundance of RNP-associated small RNAs relative to total small RNAs, corresponding to scatterplots in (D). Boxes indicate 75_th_ and 25_th_ percentiles, horizontal thick lines denote medians, and whiskers indicate the maximum and minimum values. **(F)** Pairwise comparisons of spike-in-normalized abundance of RNP-associated small RNAs in mouse testes and sperm. Each point represents the mean spike-in-normalized abundance of small RNAs derived from one locus across biological replicates. A pseudo-count of 0.5 was added before log transformation. The solid and dashed lines denote the identity line and ±10-fold fold-change line, respectively. **(G)** Ratios of the mean abundance of RNP-associated relative to total small RNAs, corresponding to scatterplots in (F). Boxes indicate 75_th_ and 25_th_ percentiles, horizontal thick lines denote medians, and whiskers sindicate the maximum and minimum values.

To test these hypotheses, we isolated ribonucleoproteins (RNPs) from mature sperm using an anion-exchange column and analyzed associated small RNAs by sequencing (TraPR) (Grentzinger *et al*, 2020) (**Fig. 4B**). As a control, capture of MIWI, MILI, and other known spermatogenic RNA-binding proteins—such as YBX1, YBX2, and YBX3 (Matsumoto & Wolffe, 1998)—in TraPR RNPs from adult mouse testes was confirmed by label-free mass spectrometry analyses (**Figs. S4A–S4B** and **Table 4**). Among 1,831 consistently detected proteins across replicate RNP samples (77.8% of all proteins detected across all replicates), 1,149 (62.8% of consistently detected proteins) were annotated RBPs (RBP2GO) (Caudron-Herger *et al*, 2021). This estimation was above the ∼7.5–9.5% of the mouse proteome identified as RBPs (Gerstberger *et al*, 2014; Hentze *et al*, 2018; Liao *et al*, 2025), confirming RBP enrichment in control testis samples. In sperm RNPs, 576 of the 1,105 proteins (52.1%) consistently detected across biological replicates were annotated RBPs, of which 351 (60.9%) were also found in testis TraPR RNPs. Despite detection of MIWI in whole sperm lysates by TMT mass spectrometry, MIWI was undetected in sperm TraPR RNPs by label-free mass spectrometry. We also noted that sperm lysis was less efficient with the buffer used in this experimental procedure and the associated lower protein yield than that of testis samples (**Figs. S4C–S4D**). These observations suggested quantitative limitations of MIWI detection due to low protein abundance. However, they do not exclude the possibility of dissociated MIWI and piRNAs in sperm.

We observed a low read alignment rate in the TraPR RNP-associated small RNA dataset—presumably due to the low-input nature of these samples—but consistent small RNA profiles across replicates (**Fig. S4E** and **Table 1**). Pachytene piRNAs were enriched in sperm RNP fractions relative to the total sperm small RNA pool with a concurrent depletion of tRFs, mtRNA fragments, and mRNA fragments regardless of the capacitation status (**Fig. 4C**). This pattern contrasted with the testis RNP control, which preserved the already dominant MIWI/MILI-associated piRNAs and AGO-associated miRNAs in the total RNA pool. Identified RNP-associated pachytene piRNAs displayed characteristic length distribution and 5′-uridine bias, indicating their integrity (**Figs. S4F–S4G**). Furthermore, the same ten loci contributed to more than half of the detected sequences across testis and sperm RNP samples, implying that similar repertoires of pachytene piRNA species remained protein-associated throughout spermiogenesis and capacitation (**Fig. S4H**). Together, the results suggested the technical limits of MIWI detection in sperm RNPs by mass spectrometry rather than dissociation of MIWI and piRNAs in sperm cells. Pachytene piRNAs, but not other small RNAs, are enriched in testis, uncapacitated sperm, and capacitated sperm, indicating continued protein association in spermiogenesis and capacitation.

Examination of small RNA species produced from individual loci showed that total testicular pachytene piRNAs, miRNAs, tRFs, and rRNA fragments maintained their abundance in testicular TraPR RNPs, indicating their association with proteins in spermatogenesis (**Figs. 4D**–**4E**). In comparison, sperm-borne RNPs harbored pachytene piRNAs and some miRNAs but lost most of the other classes. Although both are strongly Argonaute-bound in testes, a larger portion of miRNAs dissociated from proteins in sperm (**Figs. 4F**–**4G**, left). This observation is consistent with the absence of AGO proteins in our sperm proteomic datasets and contrasts with the stable piRNA-RNP association, which persists from spermatogenesis to mature sperm, demonstrated by the high correlation between testis and sperm RNP samples. Moreover, sperm capacitation minimally affected the protein-RNA association patterns (**Figs. 4F**–**4G**, right). Our findings indicate that pachytene piRNAs remain more strongly associated with proteins during spermiogenesis, sperm’s epididymal maturation, and capacitation compared to other RNA classes.

### Sperm-borne pachytene piRNAs are loaded onto MIWI

To determine whether pachytene piRNAs are indeed loaded onto MIWI in sperm, we isolated MIWI RNPs from uncapacitated sperm of *StrepII-Miwi^ki/ki^* mice, which expressed endogenously StrepII-tagged MIWI (**Fig. 5A**) (Raad *et al*, 2026). Because MIWI is far more abundant in adult testes, we first validated StrepII-MIWI purification using *StrepII-Miwi^ki/ki^* adult testis lysates. StrepII-MIWI was efficiently enriched in eluates from *StrepII-Miwi^ki/ki^*testes, but not from *StrepII-Miwi^+/+^* negative controls, as confirmed by Western blotting and silver staining (**Figs. S5A–S5B**). Mass spectrometry analyses of the eluates identified 377 proteins in the *StrepII-Miwi^ki/ki^* sample, of which 270 (71.6% of total detected proteins) were absent from the negative control (**Fig. S5C** and **Table 5**). The MIWI protein (UniProt ID: Q9JMB7) was identified with 141 peptide-spectrum matches and 43 unique peptides, corresponding to 51% sequence coverage. Furthermore, established MIWI-interacting proteins, such as PIWIL2 (MILI), TDRD2 (TDRKH), and MOV10L1, were absent from the negative control (Arif *et al*, 2022; Chen *et al*, 2009; Kojima *et al*, 2009; Takemoto *et al*, 2016; Vagin *et al*, 2009) (**Fig. S5D**).

**Figure 5.**
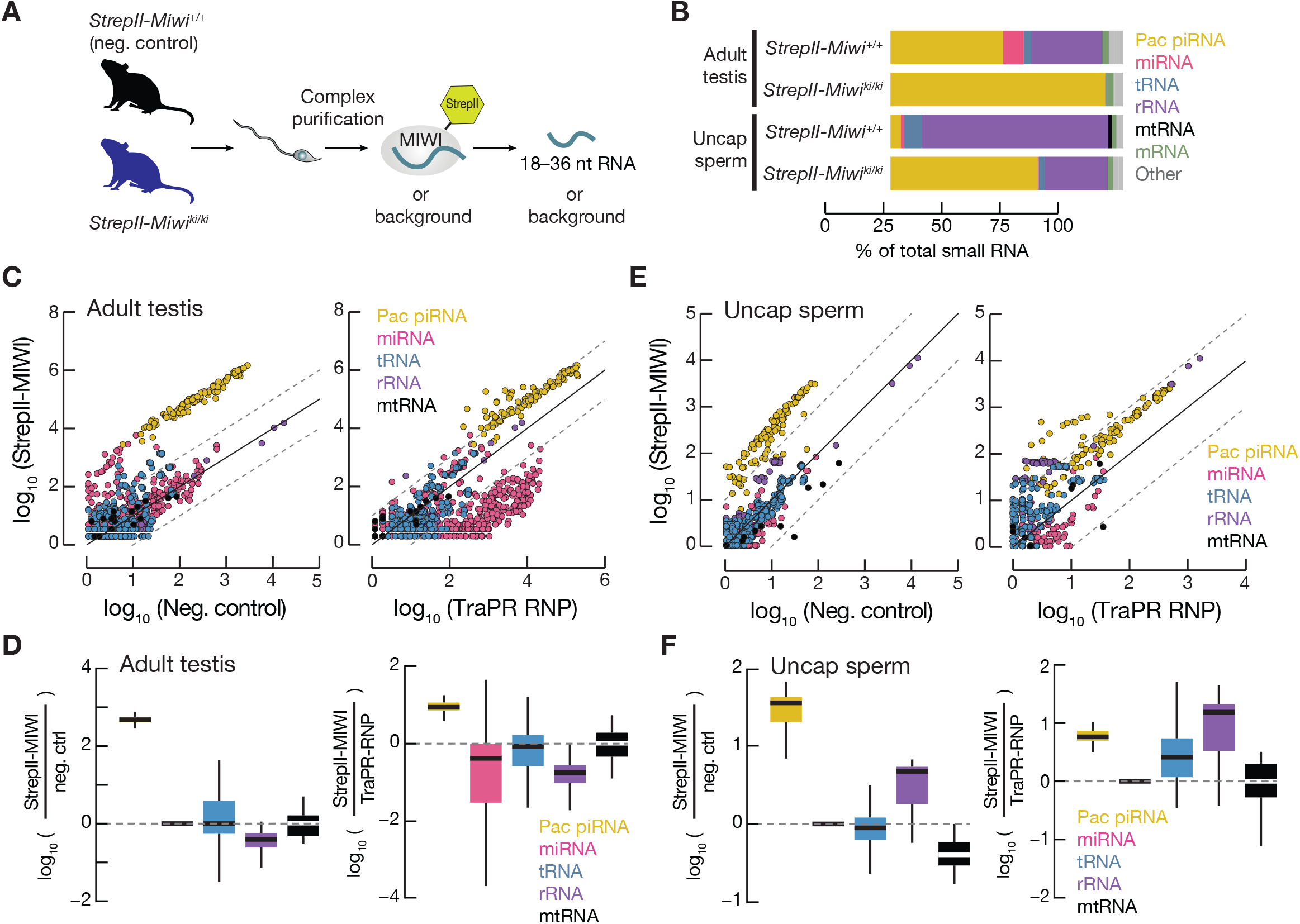
Pachytene piRNAs remain MIWI-bound in mouse sperm. **(A)** Schematic depicting StrepII-MIWI purification from adult *StrepII-Miwi^ki/ki^* mouse testes and uncapacitated cauda sperm. **(B)** Relative abundance of small RNAs identified in StrepII-MIWI eluates from adult mouse testes (*n*=1) and uncapacitated sperm (*n*=1). Each sperm sample pooled cells from five males. **(C)** Pairwise comparison of StrepII-MIWI-associated small RNAs relative to the negative control (*n*=1; left) or TraPR RNP-associated small RNAs (*n*=3; right) in adult testes. Each point represents the spike-in-normalized mean value across biological replicates. Solid and dashed lines denote the identity and ±10-fold fold-change lines, respectively. **(D)** Fold enrichment of StrepII-MIWI-associated small RNAs over the negative control (left) or TraPR RNP-associated testicular small RNAs (right), corresponding to (C). Boxes indicate the 75_th_ and 25_th_ percentiles, horizontal thick lines represent the medians, and whiskers denote the maximum and minimum values. **(E)** Pairwise comparison of StrepII-MIWI-associated small RNAs relative to the negative control (left) or TraPR RNP-associated small RNAs (*n*=3; right) in uncapacitated sperm. Each point represents the spike-in-normalized mean value of biological replicates. Solid and dashed lines denote the identity and ±10-fold fold-change lines, respectively. **(F)** Fold enrichment of StrepII-MIWI-associated sperm small RNAs over the negative control (left) or TraPR RNP-associated small RNAs (right), corresponding to (E). Each point shows the spike-in-normalized mean value of small RNAs derived from one locus. Boxes indicate 75_th_ and 25_th_ percentiles, horizontal thick lines denote medians, and whiskers indicate the maximum and minimum values.

Consistent with successful isolation of testicular StrepII-MIWI and its interacting partners, pachytene piRNAs dominated the associated small RNA pool (**Fig. 5B** and **Figs. 5C–D**, left). In contrast, AGO-associated miRNAs, which were readily detectable in TraPR RNPs, were depleted from StrepII-MIWI RNPs in testes (**Figs. 5C–D**, right), demonstrating specific isolation of endogenous StrepII-MIWI RNPs from *StrepII-Miwi^ki/ki^*cells. We employed the same procedure to purify and analyze StrepII-MIWI-associated small RNAs from *StrepII-Miwi^ki/ki^* uncapacitated sperm. Similar to testes, the StrepII-MIWI-associated small RNA pool in sperm comprised mostly pachytene piRNAs (**Fig. 5B** and **Figs. 5E–F**, left). Importantly, pachytene piRNA repertoires associated with StrepII-MIWI and TraPR RNPs were well correlated in testes and sperm, respectively. The correlation was particularly pronounced for the most abundant pachytene piRNA pools produced by distinct loci, demonstrated by similar piRNA profiles across individual loci between StrepII-MIWI and TraPR samples (an example is shown in **Fig. S5E)** and similar fractions of pachytene piRNAs derived from the same loci (**Fig. S5F**). Together, these observations indicate that the two independent strategies yielded essentially the same MIWI-associated piRNA pools.

In addition to pachytene piRNAs, a noticeably larger fraction of rRNA-derived fragments was present in sperm MIWI RNPs than in testicular MIWI RNPs. However, rRNA fragments constituted large fractions in the negative controls for both testis (30.4%) and sperm (80%) StrepII-MIWI RNPs. Together with their substantially higher abundance in mature sperm than in precursor germ cells (*i.e.*, spermatids; **Fig. 1A**), this suggests that rRNA fragments isolated by this experimental procedure represent primarily nonspecific carryover rather than biologically meaningful interactions between MIWI and the fragments. The greater abundance of rRNA fragments in *StrepII-Miwi^ki/ki^* sperm eluates than in testis eluates may therefore be attributable to increased nonspecific carryover during affinity purification.

As TraPR does not distinguish different RNA-binding proteins, the method is expected to capture protein-associated small RNAs irrespective of the protein identity (Grentzinger *et al*, 2020). Indeed, testicular miRNAs were retained in TraPR RNPs but depleted in StrepII-MIWI RNPs (**Figs. 5C–D**, right). In contrast, StrepII-MIWI and TraPR RNP-associated miRNA profiles differed little in sperm (**Figs. 5E–F**, right), consistent with the prediction of few miRNA-AGO interactions in sperm due to no or low abundance of AGO proteins. These results demonstrate that sperm-borne pachytene piRNAs are associated with MIWI and their interaction is maintained in mature sperm. In comparison, sperm miRNAs exist mainly in an unbound state rather than in AGO complexes.

## Methods

### Animals

All animal experiments were conducted in accordance with Swiss federal and cantonal regulations and approved by the veterinary authorities of the Canton of Geneva (licenses GE208A and GE507). Experiments were performed using 12–16-week-old C57BL/6J males, *StrepII-Miwi^ki/ki^* males, or *StrepII-Miwi^+/+^* littermates. The *StrepII-Miwi^ki/ki^* line (Raad *et al*, 2026) was maintained on a C57BL/6J background and shared by Ramesh Pillai. All animals were housed at the SOPF animal facility at the University of Geneva, Faculty of Medicine, and euthanized in accordance with approved protocols.

### Mouse sperm collection and purification

Sperm cells were collected from the mouse cauda epididymis. Dissected cauda tissues were rinsed in 1× PBS and transferred to pre-warmed EmbryoMax human tubal fluid (HTF) medium that contains bovine serum albumin and calcium (Sigma-Aldrich, St. Louis, MO, USA, cat. no. MR-101-D). Small incisions were made to release sperm, followed by incubation at 37°C with 5% CO_2_ for 10 min or 90 min to obtain uncapacitated and capacitated sperm, respectively (Boitrelle *et al*, 2021; Quinn *et al*, 1985; Wu *et al*, 2020). Released sperm were collected by centrifugation at 2,000 × g for 5 min at 4°C and washed with cold 1× PBS. Unwanted somatic cells were removed by purifying the collected sperm on a two-step Percoll gradient (45% and 90%; Cytiva, Marlborough, MA, USA, cat. no. 17-0891-02) followed by centrifugation at 650 × g for 25 min at 4°C. We opted for this method instead of using a detergent-containing somatic cell lysis buffer to preserve sperm membrane and RNA profile (Johnson *et al*, 2015; Pearson *et al*, 2018). Sperm at the interface were collected, diluted in excess cold PBS, and pelleted by centrifugation. Sperm purity was assessed by light microscopy and RNA sequencing. Following a final wash in cold PBS, pellets were lysed in lysis buffer and processed immediately for RNA extraction or flash-frozen for storage.

### Assessment of sperm cell lysis

Sperm collected from one mouse were divided into two equal fractions. One fraction was incubated in mirVana (Invitrogen, Waltham, MA, USA, cat. no. AM1561) or TraPR (Lexogen GmbH, Vienna, Austria, cat. no. 128.24) (Grentzinger *et al*, 2020) lysis buffer, while the control fractions were incubated in an equal volume of 1× PBS. Samples were incubated on ice for 30 min, and aliquots were mounted on glass slides with coverslips and imaged using bright-field microscopy (Zeiss Axio Imager M2) at 40× magnification.

### RNA extraction from mouse sperm

RNA extraction was performed using the mirVana miRNA Isolation Kit (Invitrogen, Waltham, MA, USA, cat. no. AM1561) following the manufacturer’s instructions for isolation of >17 nt total RNA with modifications. Cauda sperm collected from mice were lysed in lysis buffer on ice with five passages through a 27G needle using a syringe, followed by a 30 min incubation on ice after adding miRNA homogenate buffer. RNA was purified using phenol/chloroform (pH 4.3) and a mirVana column and further concentrated using RNA extraction with phenol/chloroform (pH 4.3) and ethanol precipitation.

### Analyses of sperm transcriptome by SMART-seq

Total RNA was extracted from uncapacitated C57BL/6 cauda sperm as described above. Three technical replicate libraries were constructed from 100, 250, and 500 ng of total RNA using the Takara SMART-Seq v4 ultra-low input RNA Kit (Takara Bio USA, Inc., San Jose, CA, USA, cat. no. 634890) according to the manufacturer’s instructions and were sequenced with 100 bp paired-end reads on an Illumina NovaSeq 6000 at the University of Geneva Genomics Platform. Sperm SMART-seq and published spermatogenic cell RNA-seq datasets (Wu *et al*, 2020) were processed in parallel using a standard RNA-seq pipeline with modifications to accommodate more fragmented RNAs in sperm. Briefly, adapter sequences were soft-clipped from raw reads. The resulting reads were aligned to the mouse reference genome (mm39) using STAR v2.7.11b (Dobin *et al*, 2013) with the parameter --outFilterMatchNminOverLread 0.3. Exonic reads were quantified using HTSeq v0.11.3 (Anders *et al*, 2015) and GENCODE gene annotations. Gene expression was normalized to sequencing depth with a pseudo-count of 0.5 added before log transformation. Boxplots were generated using R v4.6.1. Marker genes for sperm and cauda epididymis were selected based on published intact sperm mRNAs (Sun *et al*, 2021) and the caudal epididymal transcriptome (Rinaldi *et al*, 2020).

### Small RNA-seq library construction and data analysis

Small RNA-seq libraries were generated as described previously (Wu *et al*, 2020; Fu *et al*, 2018). Unless otherwise noted, each biological replicate refers to sperm collected from one mouse. Briefly, 1 fmol of a synthetic spike-in mixture containing 22-, 26-, and 30-nt RNA oligonucleotides (**Supplementary Table 1**) was added to 1 µg of total RNA before size selection of 18–36 nt small RNA on a 15% denaturing polyacrylamide gel. Purified small RNAs were ligated to a 3′ adapter, purified on a 10% denaturing polyacrylamide gel, and subsequently ligated to 5′ adapters containing unique molecular identifiers (UMIs). Following cDNA synthesis, libraries were PCR-amplified and purified on a 2.5% agarose gel. Library quality was assessed using a TapeStation 4150 (Agilent Technologies, Santa Clara, CA, USA). Sequencing was performed on an Illumina NovaSeq 6000 in 50-bp single-end mode at the Genomics Platform at the University of Geneva.

For data processing, adapter sequences and PCR duplicates were first removed from raw sequencing reads using Cutadapt v4.2 (Martin, 2011) and UMI-tools (Fu *et al*, 2018), respectively. Reads were then aligned using Bowtie v 1.3.1 (Langmead *et al*, 2009) and quantified with HTSeq v0.11.3 (Anders *et al*, 2015). For restricted read alignment (**Fig. 1**), deduplicated reads were sequentially aligned to annotation references in a hierarchical order: spike-ins (-v 0 -a -m 1 --best --strata), rRNA (NCBI RefSeq rRNA sequences, *Mus musculus*, GRCm39; -v 1 -a -m 3 --best --strata), tRNA (Chan & Lowe, 2016) (GtRNAdb high confidence tRNA gene set; -v 2 -a -m 3 --best --strata), miRNA (Kozomara *et al*, 2019) (miRBase; -v 2 -a -m 1 --best --strata), piRNA-producing loci (Li *et al*, 2013) (-v 2 -a -m 1 --best --strata), and the mouse genome (mm39; -v 1 -a -m 1 --best --strata). Unaligned reads from each step were carried forward to the next alignment step. For permissive alignment of all mappers (**Figs. 2**–**5**), Bowtie parameters -v 2 -k 1 - -best were used while maintaining the same hierarchical read assignment. The published spermatid small RNA-seq dataset (Wu *et al*, 2020) in Fig. 1 was generated using an identical protocol and processed in parallel using the same parameters. Read abundance was normalized using spike-in–based size factors calculated with DESeq2 (Love *et al*, 2014) unless otherwise specified. Differential expression analysis was performed using DESeq2.

### Purification of ribonucleoprotein-associated small RNAs

RNPs were purified from C57BL/6 cauda sperm using TraPR Small RNA Isolation kit (Lexogen GmbH, Vienna, Austria, cat. no. 128.24) (Grentzinger *et al*, 2020) following the manufacturer’s instructions with modifications. Briefly, sperm from two mice were lysed in 300 µl of lysis buffer on ice with five passages through a 27G needle using a syringe. After clarifying the lysate by centrifugation, the samples were loaded onto the anion-exchange resin and incubated at 18°C for 30 min. RNPs were eluted by centrifugation, and associated RNA was purified using phenol/chloroform (pH 4.3) and ethanol precipitation. 18–36 nt RNA was subsequently isolated by size selection on a 15% denaturing polyacrylamide gel. Spike-in oligos were added to each sample at 1 fmol per 1 µg of RNA before small RNA-seq library construction.

### StrepII-MIWI ribonucleoprotein complex isolation

Cauda sperm were collected from five *StrepII-Miwi*^ki/ki^ or *StrepII-Miwi*^+/+^ (negative control) mice and purified using a 27% Percoll gradient as previously described (Nixon *et al*, 2015). Sperm cells were lysed in a buffer containing 25 mM Tris-HCl (pH 8.0), 150 mM NaCl, 1 mM EDTA, 0.2% NP-40 (v/v), 0.5% sodium deoxycholate (w/v), 5% glycerol (v/v), and EDTA-free protease inhibitors (Roche, Basel, Switzerland, cat. no. 04693132001). The cleared lysates were collected by centrifugation at 16,000 × g for 15 min at 4°C for StrepII-MIWI protein complex purification using Strep-Tactin XT gravity flow columns (IBA Lifesciences GmbH, Göttingen, Germany, cat no. 2-5013-001) following the manufacturer’s instructions. As a control, testes from the same animals were processed in parallel. StrepII-MIWI purification was verified by Western blotting using rabbit polyclonal anti-PIWIL1 (MIWI) antibodies (Abcam, Cambridge, UK, cat. no ab12337; **Supplementary Table 2**). For sequencing of StrepII-MIWI-associated small RNAs, eluted proteins were concentrated using a Amicon Ultra centrifugal filter with a 10 kDa molecular weight cutoff (Milipore, Burlington, MA, USA, cat. no. UFC501024) and treated with proteinase K, followed by RNA extraction using phenol/chloroform (pH 4.3) and ethanol precipitation. Spike-in oligos were added to each sample at 1 fmol per 1 µg of RNA before small RNA-seq library construction following the procedure described above.

### Western blotting

Total sperm protein lysates were prepared as previously described (Baker *et al*, 2006). Western blotting was performed using standard procedures. Briefly, sperm samples (one mouse per replicate) were incubated in 400 µl lysis buffer containing 50 mM Tris-HCl (pH 8.5), 4% CHAPS (w/v), 7 M urea, and 2 M thiourea for 2 h on ice with vortexing every 10 min. Lysates were cleared by centrifugation at 13,000 × g for 10 min at 4°C, and supernatants were collected and concentrated using a Amicon ultra centrifugal filter with a 50 kDa molecular weight cutoff (Merck, Darmstadt, Germany, cat. no. UFC5050). Twenty percent of the final volume was resolved on a 4–20% mPAGE Bis-Tris gel (Millipore, Burlington, MA, USA, cat. no. MP42G10). After membrane transfer and blocking in fluorescent Western blot blocking buffer (Rockland Immunochemicals, Pottstown, PA, USA, cat. no. MB-070-003), tyrosine phosphorylation was detected by incubation with mouse monoclonal anti-phospho-tyrosine antibody (1:1000; Cell Signaling Technology, Danvers, MA, USA, cat. no. 96215) and polyclonal rabbit anti-TUBA4A antibody (1:5000; Abcam, Cambridge, UK, cat. no. 228701) overnight at 4°C. After washing with TBS with 0.1% Tween-20 (v/v), membranes were incubated with IRDYE-conjugated donkey anti-mouse and goat anti-rabbit antibodies (1:10,000; LI-COR Biosciences, Lincoln, NE, USA, cat. nos. 926-68072 and 926-32211, respectively). Fluorescent signals were detected using the LI-COR Odyssey imaging system (LI-COR Biosciences, Lincoln, NE, USA). See also **Supplementary Table 2.**

### Label-free mass spectrometry

For analyses of TraPR- and StrepII-MIWI eluates, proteins were digested with a mix of Trypsin/LysC using the iST kit (PreOmics GmbH, Planegg/Martinsried, Germany), and peptides were analyzed by nanoLC-ESI-MSMS using an easy-nLC 1000 liquid chromatography system (Thermo Fisher Scientific, Waltham, MA, USA) coupled with a Q-Exactive HF Hybrid Quadrupole-Orbitrap Mass Spectrometer at the Proteomics Platform at the University of Geneva. Database searches were performed with Mascot (Matrix Science Ltd, London, UK) using the mouse reference proteome database (UniProt) (The UniProt Consortium *et al*, 2025) as well as the sequence of the expressed MIWI protein (UniProt ID: Q9JMB7). Data were analyzed and validated with Scaffold v5.3.3 (Proteome Software Inc., Portland, OR). Peptide identifications were accepted if they could be established at greater than 57% probability to achieve an FDR less than 0.001 by the Percolator posterior error probability calculation (Käll *et al*, 2008). Protein identifications were accepted if they could be established at greater than 99% probability to achieve an < 0.01 FDR and contained ≥ 2 identified peptides. Protein probabilities were assigned by ProteinProphet (Nesvizhskii *et al*, 2003). Proteins sharing identified peptides and therefore indistinguishable by MS/MS analysis were grouped according to the principle of parsimony.

### Tandem mass tag mass (TMT) spectrometry

C57BL/6 cauda sperm were lysed in 20 mM HEPES (pH 8.0), 8 M urea, EDTA-free protease inhibitors (Roche, Basel, Switzerland, cat. no. 04693132001), and phosphatase inhibitor cocktail 3 (Sigma-Aldrich, St. Louis, MO, USA, cat. no P0044). Samples were sonicated using a Branson Digital Sonifier 450 (Branson Ultrasonics Corp., Danbury, CT, USA) with 10% amplitude. Lysates were cleared by centrifugation at 20,000 × g for 15 min at room temperature. FACS-sorted spermatids were lysed using a two-step procedure. Urea lysis buffer was added to the cells, followed by sonication using a Branson Sonifier at 80% amplitude in an ice-water bath for 10 cycles of 12 s each, with 45 s cooling intervals on ice between cycles. Lysates were cleared by centrifugation at 20,000 × g for 15 min at 4°C. After the supernatant was transferred to fresh tubes, RIPA buffer containing 25 mM Tris-HCl (pH 7.5), 150 mM NaCl, 1% NP-40 (v/v), 1% sodium deoxycholate (w/v), 0.1% SDS (w/v), protease inhibitors, and phosphatase inhibitors was added to the cell pellet. A Covaris ultrasonicator (Covaris, Woburn, MA, USA) was used for a second round of sonication under the following conditions: 1,200 s total processing time, 10% duty factor, 175 W peak incident power, and 200 cycles per burst. Lysates were cleared by centrifugation at 20,000 × g for 15 min at room temperature. The supernatant was collected and combined with the first supernatant and flash-frozen until further processing. Protein concentrations were measured using the BCA assay (Thermo Fisher Scientific, Waltham, MA, USA, cat no. 23225), and TMT mass spectrometry analyses were performed at the Proteomics Resource Center at New York University (New York, USA).

Proteins were digested with trypsin, and peptides were labeled with TMT10plex isobaric label reagent (Thermo Fisher Scientific, Waltham, MA, USA, cat no. 90309). Combined samples were fractionated off-line by basic reverse-phase HPLC, and 15 fractions were collected. Each fraction was analyzed using a Q Exactive Orbitrap mass spectrometer (Thermo Fisher Scientific, Waltham, MA, USA), and the data were searched against the mouse reference proteome database using the Andromeda search engine in MaxQuant (Tyanova *et al*, 2016). Data were filtered to achieve an FDR < 0.01 at both the peptide and protein levels using a target-decoy approach. Proteins identified by ≥ 2 unique peptides and detected in at least three out of five biological replicates were retained for downstream analysis. For quantitative comparison, a z-score was calculated for each protein using its mean TMT intensity relative to the mean intensities of all other proteins detected within the same experiment. The results, therefore, reflect the relative abundance of each protein within its respective proteome. Proteins detected in both spermatid and sperm datasets in **Fig. 4A** were assigned z-scores, enabling a comparison of relative abundance between the cell types.

## Discussion

The sperm small RNA repertoire comprises remnants of small RNA biogenesis in spermatids, including both intact sequences and degraded products. Among cytoplasmic small RNAs, piRNAs exhibit the most substantial reduction in sperm. Retained piRNAs, however, remain as a major population of small RNAs in sperm and are minimally affected by epididymal maturation (Sharma *et al*, 2016, 2018; Wang *et al*, 2023) and capacitation (**Fig. 3D**). These observations indicate that sperm-borne piRNAs are of mostly germline and testicular origin, showing no apparent newly produced (Hutcheon *et al*, 2017) or acquired species after spermatogenesis is complete. This contrasts with sperm-borne miRNAs and tRFs, for which additional species were observed after spermiogenesis (**Fig. 1B**) and the composition has been shown to be malleable after sperm formation. One proposed mechanism is *via* exchange between the sperm cytoplasm and cytoplasmic droplets, the latter contain residual cytoplasmic content and remain attached to mature sperm after cytoplasmic elimination (Wang *et al*, 2023). Another mechanism is through delivery of small RNA sequences by epididymosomes secreted by the epididymal epithelium (Sharma *et al*, 2018).

Reported composition, origin, and intergenerational effects of sperm-borne RNAs vary, which has been attributed to differences in the mouse strain, sperm sample preparation method, or procedure of proof-of-function experiments (Chen *et al*, 2016; Conine *et al*, 2020; Hutcheon *et al*, 2017; Johnson *et al*, 2015; Peng *et al*, 2012; Sellem *et al*, 2020; Sharma *et al*, 2016; Short *et al*, 2017; Stanger *et al*, 2020; Tan *et al*, 2020; Tomar *et al*, 2024; Wang *et al*, 2023, 2020). Technical differences in sequencing analyses, including the size selection range for library construction and read alignment strategy, may further contribute to discrepancies. Moreover, absolute quantification of mouse sperm-borne small RNAs reveals supra-physiological concentrations of RNAs commonly used in proof-of-function studies of paternally inherited small RNAs (König *et al*, 2025). These observations underscore the need for a more quantitative and physiologically informed understanding of sperm-borne small RNAs. Despite the ideal experimental setup for absolute quantification of RNAs in this study, the variability and imprecision in sperm counts obtained by hemocytometry among technical replicates and experimenters in our hands and described in published reports (Brazil *et al*, 2004; Prathalingam *et al*, 2006; Tomlinson *et al*, 2001) were incompatible with the level of precision required for absolute per-cell RNA quantification. An unbiased and reproducible sperm-counting method will facilitate accurate absolute quantification of sperm-borne RNAs in future studies.

Here, we identified two additional confounding factors associated with the transcriptionally inert nature of sperm and low RNA yield common to sperm sample preparation. Firstly, RNA decay in sperm extends beyond rRNAs and spermiogenesis, progressively fragmenting functional spermatid rRNAs, small RNAs, and mRNAs until at least capacitation (**Figs. 1** and **3**). For example, pre-pachytene piRNAs, whose expression peaks before the pachytene stage of meiosis in spermatogenesis (Aravin *et al*, 2007, 2008; Li *et al*, 2013; Ozata *et al*, 2019), can be detected in cauda sperm but mostly as degraded products. rRNA decay is already pronounced in uncapacitated cauda sperm, whereas degradation of mRNAs encoding ribosomal proteins escalates at capacitation. Whether this observation reflects differences in intrinsic RNA stability or a regulated, step-wise process to repress translation in mature sperm (Johnson *et al*, 2011; Lei *et al*, 2021) is unclear. Verifying intact species is straightforward for Argonaute-bound small RNAs with experimentally established lengths, nucleotide biases, and characteristic RNA modifications (*i.e.*, 3′ 2′-*O*-methylation of mouse piRNAs) (Bartel, 2018; Ozata *et al*, 2019) but has proven challenging for others with less defined characteristics.

Secondly, to the best of our knowledge, few published reports address PCR duplicates that often inadvertently accompany low-input sperm small RNA sequencing samples (Conine *et al*, 2018; Gòdia *et al*, 2018). UMI incorporation remains a gold standard for distinguishing technically identical reads from biologically identical ones and improves quantification accuracy across sequencing methods (Fu *et al*, 2018; Hong & Gresham, 2017; Kivioja *et al*, 2011; Smith *et al*, 2017; Sun *et al*, 2024; Zhu *et al*, 2020). Our UMI-integrated data demonstrate that PCR duplicates inflate the estimated quantities of abundant small RNAs. tRFs are particularly prone to artifactual duplicates, likely due to the small size (74–93 nt), multi-copy genomic origin, and high abundance of the source tRNAs (Parisien *et al*, 2013). The rank order of tRFs is minimally affected, but their abundance may be overestimated in the absence of duplicate removal. Importantly, the most susceptible tRF species (*e.g*., 5′ fragments derived from *tRNA-Gly-GCC* and *tRNA-Glu-CTC*) are also most sensitive to capacitation (**Fig. 3D**) and have been identified as epididymosome-delivered, paternally inherited species in embryos (Chen *et al*, 2016; Sharma *et al*, 2016, 2018). In contrast to tRFs, pachytene piRNAs are produced from large, repeat-poor genomic loci (median precursor transcript length: ∼31,826 nt) (Li *et al*, 2013; Yu *et al*, 2021). They produce far fewer PCR duplicates under the same experimental conditions (**Fig. S2D**) and show higher concordance between independently generated datasets regardless of UMI-based deduplication (**Fig. 2D**). A limitation of our sequencing strategy, as well as those used in other reports, is the selective capture of RNAs bearing 5′-monophosphate and 3′-hydroxyl termini. As tRFs could harbor 3′ terminal 2′,3′-cyclic phosphates, dense RNA modifications, and stable secondary structures (Padhiar *et al*, 2024), dedicated methods (Gu *et al*, 2022; Gustafsson *et al*, 2025; Shi *et al*, 2021b) combined with UMIs would be necessary to accurately quantify sperm tRFs.

Using UMI-based deduplication and spike-in-based normalization, we observed reproducible small RNA profiles across biological replicates. This could in theory be explained by the isogenic background of C57BL/6 mice alone. However, inbred mice do not necessarily exhibit uniform fertility traits (Sztein *et al*, 2000; Tuttle *et al*, 2018), suggesting that selective RNA elimination or retention may play a role in forming a consistent sperm small RNA profile. Cytoplasmic elimination may represent a window of opportunity, as this evolutionarily conserved process is orderly, essential for male fertility, and strictly regulated by SPEM1, ARRDC5, and TEX38 (O’Donnell *et al*, 2011; Rengan *et al*, 2012; Sprando & Russell, 1987; Yang *et al*, 2026; Zheng *et al*, 2007). Another window of opportunity for sperm RNA content remodeling during the epididymal transit has been described (Sharma *et al*, 2016, 2018; Wang *et al*, 2023). How RNA selectivity may be reinforced by cytoplasmic elimination, epididymosome-mediated RNA acquisition, or cytoplasmic droplet-mediated RNA exchange remains unknown. A possibility is a yet-to-be-identified, sperm-specific cytoplasmic RNP structure akin to chromatoid bodies in developing mammalian spermatogenic cells (Kotaja & Sassone-Corsi, 2007; Meikar *et al*, 2014; Schreier *et al*, 2022). Alternatively, subcellular regions within the highly compact and structured sperm cell may actively retain or inadvertently entrap specific RNA species (Johnson *et al*, 2015; Korneev *et al*, 2021; Mortimer, 2018). For example, although our data support sperm cytoplasm as the primary site of RNA remodeling (**Fig. 1E**), we cannot exclude the possibility that the presumed cytoplasmic small RNAs in our analyses become entrapped in the nucleus during chromatin compaction at spermiogenesis.

The small RNA composition in sperm remains dynamic beyond epididymal maturation. Artificially induced capacitation further elevates subsets of individual tRFs and miRNAs (**Fig. 3D**), suggesting a potential for continued refinement of the RNA repertoire immediately before fertilization *in vivo*. These observations have implications for paternal inheritance studies and assisted reproductive procedures (*i.e.*, intracytoplasmic sperm injection), for which uncapacitated sperm are routinely used. Questions about the long-term consequences of using uncapacitated sperm and spermatids to achieve fertilization have been raised but remain unresolved (Esteves *et al*, 2018). Among candidate paternal epigenetic molecules delivered to embryos only under *in vivo* conditions, how capacitation-sensitive small RNAs may influence embryo viability has yet to be determined. Our sperm capacitation experiment adopted an established combination of chemicals and *in vitro* incubation conditions (see Methods) that reproduces many molecular hallmarks of physiological capacitation. However, it does not entirely recapitulate the *in vivo* environment (Boitrelle *et al*, 2021). Future experiments using ZP3 recombinant proteins or progesterone (Beebe *et al*, 1992; Brewis *et al*, 1996; Foresta *et al*, 1992) may help confirm the physiological relevance of our observations and elucidate the source of increased tRFs and miRNAs in our capacitated sperm data.

The most abundant pachytene piRNA pools produced by distinct loci in spermatogenesis persist in sperm (**Fig. 1F**) and through capacitation (**Fig. 3E**). Our proteomics data align with the presence of MIWI proteins in mature mouse sperm (Skerrett-Byrne *et al*, 2022), and the complementary RNP purification experiments demonstrate association of piRNAs with MIWI even after sperm capacitation. An unresolved observation from these experiments is our inability to detect MIWI by label-free mass spectrometry in wild-type sperm TraPR eluates or *StrepII-Miwi^ki/ki^*sperm total lysates, despite detection of MIWI in wild-type total sperm lysates by TMT-based mass spectrometry and enrichment of authentic pachytene piRNAs in all purified RNP samples. TMT-based mass spectrometry may offer improved sensitivity for detecting low-abundance purified sperm MIWI in future experiments.

While MIWI association may help protect piRNAs from widespread RNA decay, 3′ terminal 2′-*O*-methylation of piRNAs may more directly confer RNA stability. Terminal methylation stabilizes piRNAs in mouse spermatogenic cells by blocking nuclease-mediated RNA cleavage (Gainetdinov *et al*, 2021; Lim *et al*, 2015). However, as nuclease-mediated RNA degradation pathways presumably do not operate in mature sperm, terminal 2′-*O*-methylation of piRNAs may contribute to RNA stability *via* other mechanisms conferred by the modification, such as increasing RNA resistance to alkaline hydrolysis (Motorin & Marchand, 2018). Consistent with the idea that MIWI association may play a partial or indirect role in piRNA stability in sperm, while the most abundant pachytene piRNA pools from spermatids are preferentially retained in sperm (**Fig. 1F**), detected sequences from all loci are present in sperm MIWI RNPs (**Fig. 5E**). We note that a small fraction of pachytene piRNAs appear to disassociate from sperm RNPs, compared to the nearly full engagement of the pachytene piRNA pool with their PIWI protein partners in testes (**Fig. 4E**). The explanation may be informed by our proteomics data, which revealed a major difference in piRNA pathway protein composition between testicular germ cells and sperm—whereas the former harbors a high level of both pachytene piRNA-interacting PIWI proteins, MIWI and MILI, only MIWI remains detectable in mature sperm. We currently do not know whether the observed unbound sperm-borne piRNA sequences represent MIWI- or MILI-bound species from an early stage (*i.e.*, spermiogenesis or epididymal maturation). Terminal methylations may also help explain the persistence of these unbound piRNA sequences. Unlike piRNAs, most miRNAs appear to exist in an unbound state without their AGO protein partners in sperm. How unmethylated miRNAs escape RNA decay without protein association is unclear. The distinct protein association properties of sperm-borne small RNAs suggest that sperm likely deliver piRNA-MIWI complexes and unbound miRNAs to oocytes at fertilization.

In conclusion, in addition to remodeling of sperm-borne small RNA composition during epididymal transit, our findings demonstrated that selective RNA retention, stochastic degradation, and capacitation-driven changes shape the mature sperm RNA repertoire. These are important considerations for interpreting sperm RNA biology and its post-fertilization function. The relevance of these molecular forces in the sperm of mammalian species exhibiting genetic polymorphisms and heterogeneity in qualitative and quantitative fertility traits, such as humans, awaits further investigation.

## Data Availability

Sequencing data generated in this study are available at the Gene Expression Omnibus (GSE339509). RNA-seq and small RNA-seq data from C57BL/6 spermatocytes and spermatids have been published (PRJNA634688) (Wu *et al*, 2020).

## Acknowledgements

We thank the University of Geneva Animal Facility for animal husbandry; the Animal Experimental Office for assistance with animal experimentation licenses; members of the Wu, Pillai, and Nef laboratories, Ramesh Pillai, Antoine H.F.M. Peters, and Phillip D. Zamore for helpful discussions; and Ramesh Pillai for the *StrepII-Miwi* transgenic mouse line. This study was supported by the Swiss National Science Foundation PRIMA grant (PR00P3_201535) awarded to P.-H.W.

## Author Contributions

Conceptualization, P.-H.W.; supervision, P.-H.W.; methodology, G.P., G.E.A, S.S., and P.S.; investigation, G.P., S.S., K.S., B.M.U., S.C.-P., and M.V.D.; formal analysis, G.E.A, G.P., and B.M.U; visualization, G.P., G.E.A., K.S., P.-H.W, writing—original draft, G.P. and P.-H.W.; writing—review & editing, P.-H.W.; funding acquisition, P.-H.W.

## Supplementary Figure Legends

**Figure S1.**
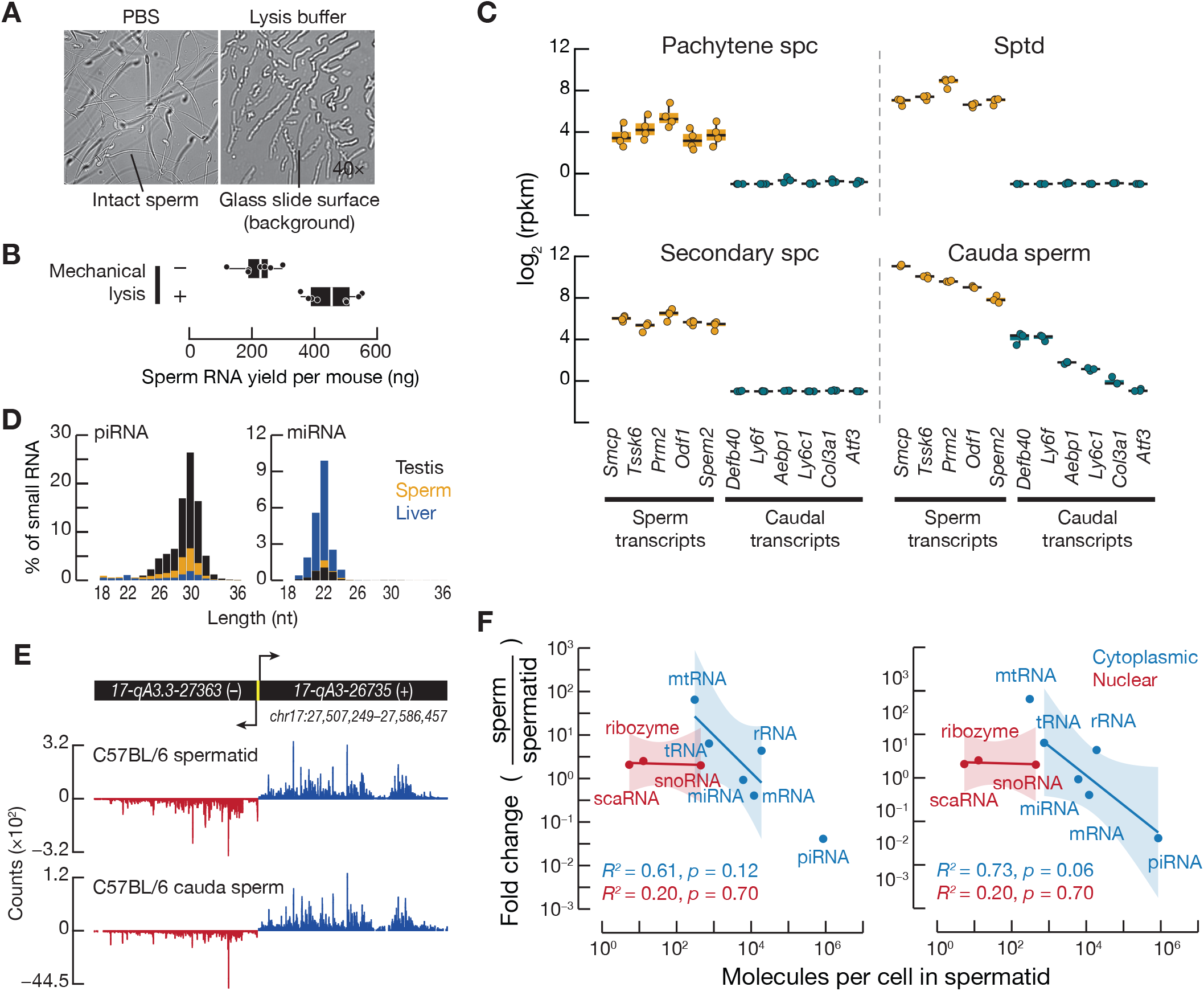
Sperm lysis efficiency and small RNA composition. **(A)** Lysis of mouse cauda sperm in PBS (control) or a commercial mirVana lysis buffer. **(B)** RNA yield from cauda sperm with or without mechanical homogenization by passing lysates through a needle using a syringe (*n*=8). Each point denotes RNA yield from purified sperm from one mouse. Boxes indicate 75_th_ and 25_th_ percentiles, vertical thick lines denote medians, and whiskers indicate the maximum and minimum values. **(C)** Expression of reported sperm-and caudal epididymis-enriched transcripts in purified mouse germ cells and cauda sperm. Each point denotes the value of a replicate sample, with a pseudo-count of 0.5 added before log transformation. Boxes indicate the interquartile range, the horizontal lines denote medians, and whiskers indicate the maximum and minimum values. Spc, spermatocyte. Sptd, spermatid. **(D)** Length distributions of detected miRNAs and piRNA sequences in C57BL/6 testis, sperm, and liver (control). **(E)** piRNA mapping profiles across the *pi17* pachytene piRNA-producing locus in spermatids and cauda sperm. The promoter is colored in yellow. **(F)** Regression analysis of the relationship between the abundance of small RNA classes in spermatids and their retention in sperm, excluding piRNA (left) or mitochondrial RNA (mtRNA) fragments (right). Each point represents the spike-in-normalized mean abundance of all non-zero RNAs in the indicated category in spermatids (*n*=3) and sperm (*n*=3). Shaded regions indicate 95% confidence intervals of the fitted regression lines. Subcellular localization assignments reflect the predominant patterns.

**Figure S2.**
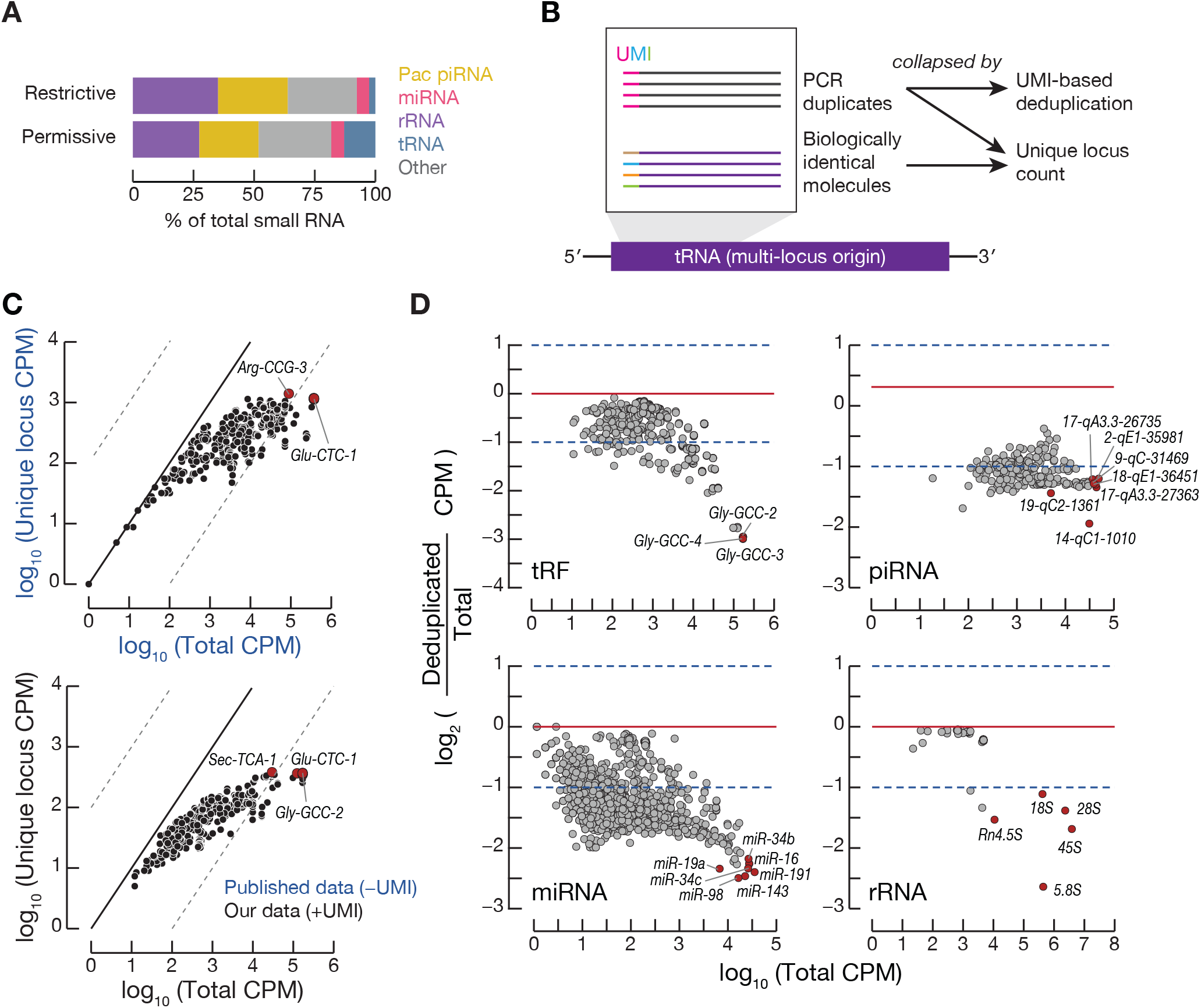
Impact of PCR duplicates on small RNA quantification in sperm. **(A)** Relative abundance of small RNAs in C57BL/6 cauda sperm. Fractions were calculated using restrictive or permissive read-alignment parameters that identify some or all multi-mapping sequences, respectively (see Methods). Pac piRNA, pachytene piRNA. **(B)** Schematics depicting UMI-based PCR duplicate removal and collapsing identical sequences regardless of origin by non-redundantly counting expressed loci. UMI, unique molecular identifier. **(C)** Estimated tRF abundance in sperm before and after collapsing all identical sequences regardless of origin. Each dot represents the mean abundance of fragments produced from a tRNA gene (*n*=3). Dashed lines indicate 100-fold differences. CPM, counts per million. **(D)** Effects of PCR duplicates on abundant tRFs, piRNAs, miRNAs, and rRNA fragments in our dataset. Each point denotes the mean value of sequences derived from a locus (*n*=3). The most abundant species are highlighted in red.

**Figure S3.**
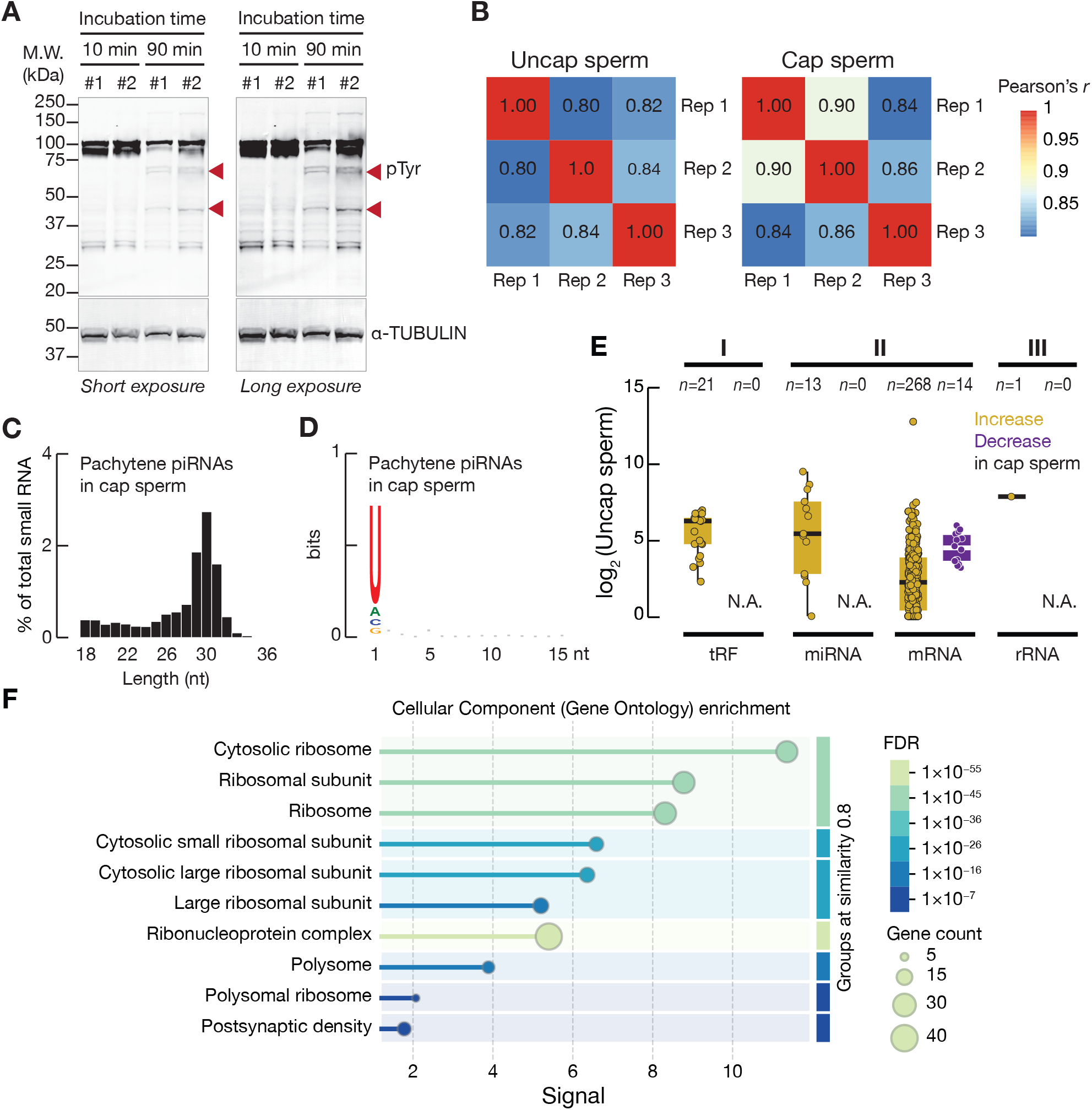
Effects of capacitation on sperm small RNAs. **(A)** Protein tyrosine phosphorylation in C57BL/6 cauda sperm incubated under capacitating conditions for 10 min (negative control) or 90 min assessed by Western blotting. Each replicate represents sperm from one mouse. Red arrowheads, increased tyrosine phosphorylation associated with sperm capacitation. pTyr, phosphorylated tyrosine. M.W., molecular weight. **(B)** Correlations among uncapacitated (uncap) or capacitated (cap) sperm samples. Calculations were performed using spike-in-normalized counts, excluding RNA species with zero reads across all samples. **(C)** Length distribution of pachytene piRNAs identified in capacitated sperm. **(D)** Nucleotide distribution of pachytene piRNAs identified in capacitated sperm analyzed using WebLogo 3.9.0 (Crooks *et al*, 2004). **(E)** Abundance of capacitation-sensitive small RNAs in (C). Boxes indicate 75_th_ and 25_th_ percentiles, horizontal thick lines denote the median, and whiskers indicate the maximum and minimum values. Each point represents the mean spike-in-normalized abundance of RNAs produced from a distinct RNA locus (*n*=3), with a pseudo-count of 1 added before log transformation. **(F)** Gene Ontology analysis of cellular components represented by proteins encoded by mRNAs with increased fragments in capacitated sperm in Fig. 3D, Category II (*n*=268). Analysis was performed using STRING v12.5 (Szklarczyk *et al*, 2025).

**Figure S4.**
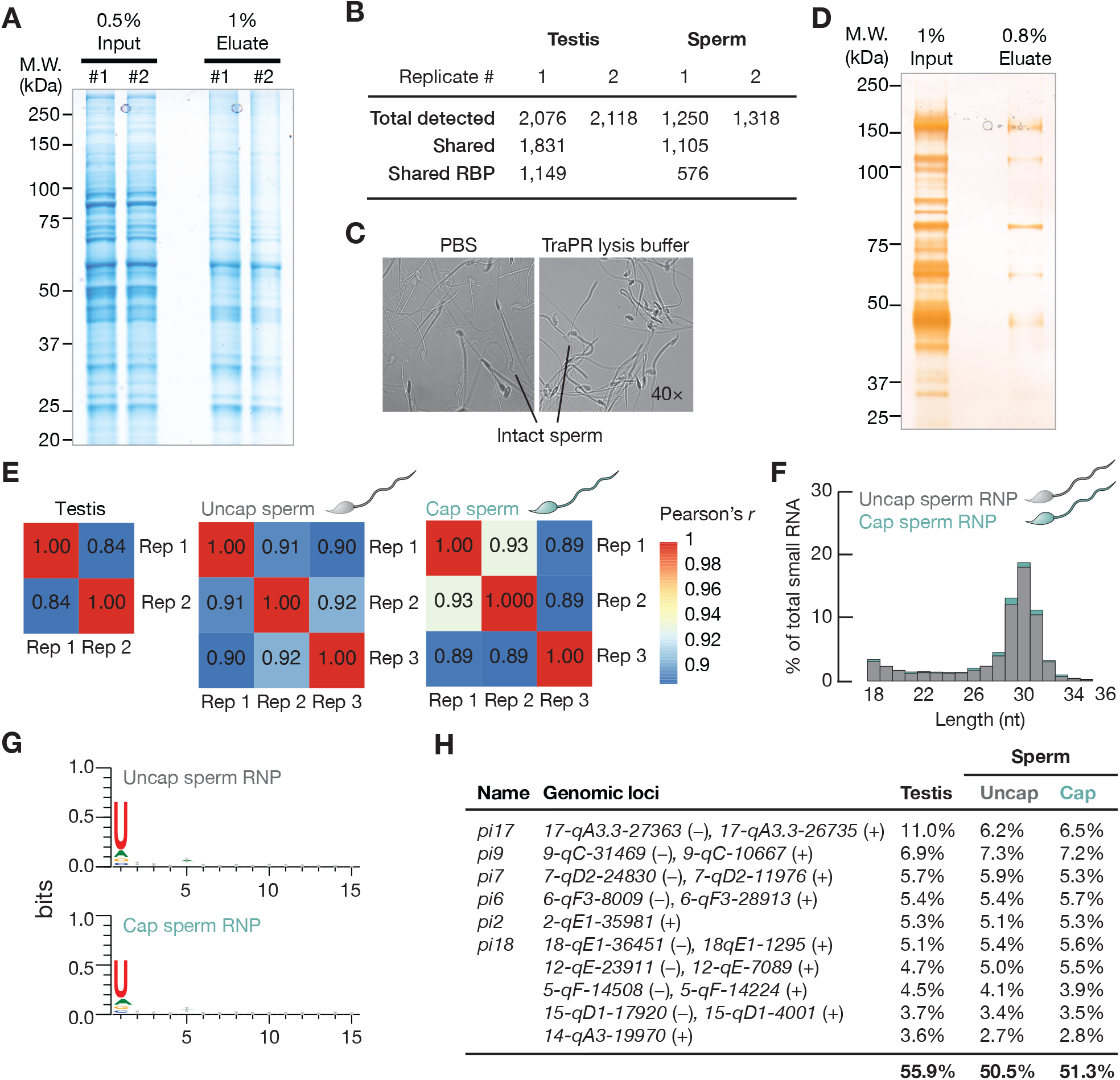
Isolation of protein-associated small RNAs from mouse testes and sperm. **(A)** Coomassie staining of gel-resolved TraPR eluates collected from adult C57BL/6 mouse testes. Replicate numbers are indicated with each replicate derived from a single mouse. M.W., molecular weight. **(B)** Summary of label-free mass spectrometry analysis of TraPR eluates from adult mouse testes or uncapacitated cauda sperm. RNA-binding proteins (RBPs) were identified using RBP2GO (Caudron-Herger *et al*, 2021). **(C)** Lysis efficiency of cauda sperm in PBS (negative control) or TraPR-compatible cell lysis buffer. **(D)** Silver staining of gel-resolved TraPR eluates from uncapacitated sperm. Each replicate sample was collected from six mice. **(E)** Pearson correlation coefficient of TraPR RNP-associated small RNA profiles in testis and sperm across replicate samples. Uncap, uncapacitated. Cap, capacitated. **(F)** Length distribution overlays of TraPR RNP-associated pachytene piRNAs in uncapacitated and capacitated sperm. **(G)** Nucleotide distributions of TraPR RNP-associated pachytene piRNAs in uncapacitated and capacitated sperm generated using WebLogo 3.9.0 (Crooks *et al*, 2004). **(H)** Genomic sources of the ten most abundant TraPR RNP-associated pachytene piRNA pools in adult testes (*n=2*) and their corresponding fractions in uncapacitated sperm (*n*=3) and capacitated sperm (*n*=3). Mean fractions of biological replicates are shown. Locus pairs sharing a bidirectional promoter are presented as a single locus. Published alternative locus names are indicated when applicable.

**Figure S5.**
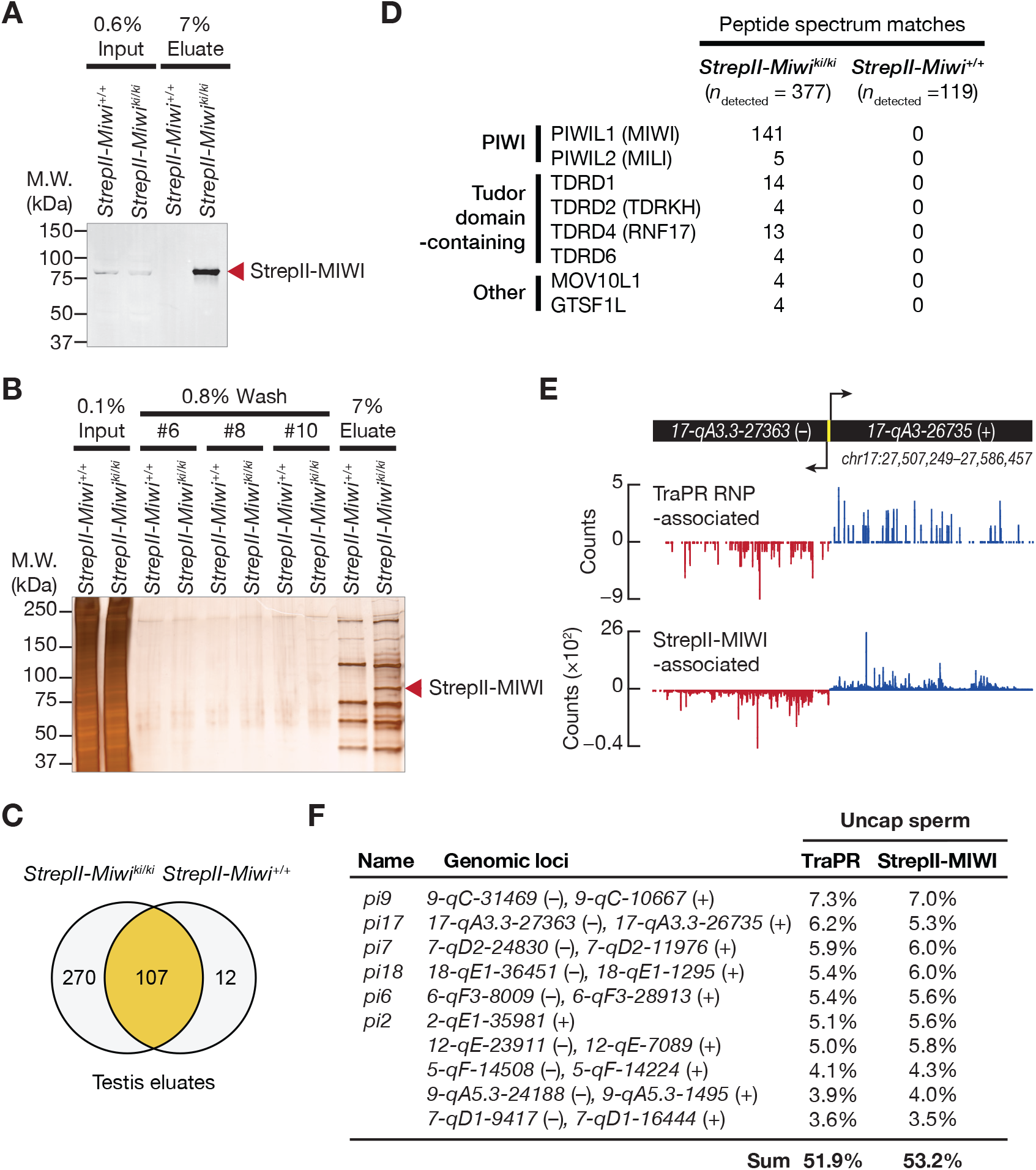
StrepII-MIWI-associated pachytene piRNAs in mouse testes and sperm. **(A)** Isolation of StrepII-MIWI from adult *StrepII-Miwi^ki/ki^* mouse testes confirmed by Western blotting. M.W., molecular weight. **(B)** Silver staining of gel-resolved StrepII-MIWI eluates collected from adult testes of *StrepII-Miwi^ki/ki^* and *StrepII-Miwi^+/+^* (negative control) mice. The band indicating StrepII-MIWI was confirmed by mass spectrometry. **(C)** Venn diagram showing proteins identified by label-free mass spectrometry in *StrepII-Miwi^ki/ki^*and *StrepII-Miwi^+/+^* testis eluates from the affinity purification. **(D)** Known MIWI-interacting piRNA pathway proteins identified in the StrepII-MIWI testis eluate. **(E)** Mapping profiles of TraPR RNP- and StrepII-MIWI-associated piRNAs across the *pi17* pachytene piRNA-producing locus. The promoter is colored in yellow. **(F)** Genomic sources of the ten most abundant pachytene piRNA pools in TraPR RNPs isolated from uncapacitated sperm (*n*=3; six mice per replicate) and their corresponding representations in StrepII-MIWI RNPs (*n*=1; five mice per replicate). Mean fractions of biological replicates are shown. Locus pairs sharing a promoter are presented as a single locus. Published alternative locus names are indicated when applicable.

## References

1. de Albuquerque BFM, Placentino M & Ketting RF (2015) Maternal piRNAs are essential for germline development following de novo establishment of endo-siRNAs in Caenorhabditis elegans. Dev Cell 34: 448–456

2. Alves MBR, De Arruda RP, De Bem THC, Florez-Rodriguez SA, Sá Filho MFD, Belleannée C, Meirelles FV, Da Silveira JC, Perecin F & Celeghini ECC (2019) Sperm-borne miR-216b modulates cell proliferation during early embryo development via K-RAS. Sci Rep 9: 10358

3. Anders S, Pyl PT & Huber W (2015) HTSeq--a Python framework to work with high-throughput sequencing data. Bioinformatics 31: 166–169

4. Aravin A, Gaidatzis D, Pfeffer S, Lagos-Quintana M, Landgraf P, Iovino N, Morris P, Brownstein MJ, Kuramochi-Miyagawa S, Nakano T, et al (2006) A novel class of small RNAs bind to MILI protein in mouse testes. Nature 442: 203–207

5. Aravin AA, Sachidanandam R, Bourc’his D, Schaefer C, Pezic D, Toth KF, Bestor T & Hannon GJ (2008) A piRNA Pathway Primed by Individual Transposons Is Linked to De Novo DNA Methylation in Mice. Mol Cell 31: 785–799

6. Aravin AA, Sachidanandam R, Girard A, Fejes-Toth K & Hannon GJ (2007) Developmentally regulated piRNA clusters implicate MILI in transposon control. Science 316: 744–747

7. Arif A, Bailey S, Izumi N, Anzelon TA, Ozata DM, Andersson C, Gainetdinov I, MacRae IJ, Tomari Y & Zamore PD (2022) GTSF1 accelerates target RNA cleavage by PIWI-clade Argonaute proteins. Nature 608: 618–625

8. Baker MA, Hetherington L & Aitken RJ (2006) Identification of SRC as a key PKA-stimulated tyrosine kinase involved in the capacitation-associated hyperactivation of murine spermatozoa. J Cell Sci 119: 3182–3192

9. Barckmann B, Pierson S, Dufourt J, Papin C, Armenise C, Port F, Grentzinger T, Chambeyron S, Baronian G, Desvignes J-P, et al (2015) Aubergine iCLIP Reveals piRNA-Dependent Decay of mRNAs Involved in Germ Cell Development in the Early Embryo. Cell Rep 12: 1205–1216

10. Bartel DP (2004) MicroRNAs: genomics, biogenesis, mechanism, and function. Cell 116: 281–297

11. Bartel DP (2018) Metazoan microRNAs. Cell 173: 20–51

12. Beebe SJ, Leyton L, Burks D, Ishikawa M, Fuerst T, Dean J & Saling P (1992) Recombinant mouse ZP3 inhibits sperm binding and induces the acrosome reaction. Dev Biol 151: 48–54

13. Bianchi E, Stermer A, Boekelheide K, Sigman M, Hall SJ, Reyes G, Dere E & Hwang K (2018) High-quality human and rat spermatozoal RNA isolation for functional genomic studies. Andrology 6: 374–383

14. Boerke A, Dieleman SJ & Gadella BM (2007) A possible role for sperm RNA in early embryo development. Theriogenology 68 Suppl 1: S147–155

15. Boitrelle F, Shah R, Saleh R, Henkel R, Kandil H, Chung E, Vogiatzi P, Zini A, Arafa M & Agarwal A (2021) The Sixth Edition of the WHO Manual for Human Semen Analysis: A Critical Review and SWOT Analysis. Life 11: 1368

16. Brazil C, Swan SH, Drobnis EZ, Liu F, Wang C, Redmon JB, Overstreet JW, & Study for Future Families Research Group (2004) Standardized Methods for Semen Evaluation in a Multicenter Research Study. J Androl 25: 635–644

17. Brennecke J, Malone CD, Aravin AA, Sachidanandam R, Stark A & Hannon GJ (2008) An epigenetic role for maternally inherited piRNAs in transposon silencing. Science 322: 1387–1392

18. Brewis IA, Clayton R, Barratt CL, Hornby DP & Moore HD (1996) Recombinant human zona pellucida glycoprotein 3 induces calcium influx and acrosome reaction in human spermatozoa. Mol Hum Reprod 2: 583–589

19. Capra E, Turri F, Lazzari B, Cremonesi P, Gliozzi TM, Fojadelli I, Stella A & Pizzi F (2017) Small RNA sequencing of cryopreserved semen from single bull revealed altered miRNAs and piRNAs expression between High- and Low-motile sperm populations. BMC Genomics 18: 14

20. Caudron-Herger M, Jansen RE, Wassmer E & Diederichs S (2021) RBP2GO: a comprehensive pan-species database on RNA-binding proteins, their interactions and functions. Nucleic Acids Res 49: D425–D436

21. Cecchini K, Zamani M, Ajaykumar N, Vega-Badillo J, Bagci A, Bailey S, Zamore PD & Gainetdinov I (2026) Cleavage of mRNAs by a minority of pachytene piRNAs improves sperm fitness. Nature 652: 508–516

22. Chan PP & Lowe TM (2016) GtRNAdb 2.0: an expanded database of transfer RNA genes identified in complete and draft genomes. Nucleic Acids Res 44: D184–D189

23. Chen C, Jin J, James DA, Adams-Cioaba MA, Park JG, Guo Y, Tenaglia E, Xu C, Gish G, Min J, et al (2009) Mouse Piwi interactome identifies binding mechanism of Tdrkh Tudor domain to arginine methylated Miwi. Proc Natl Acad Sci U S A 106: 20336–20341

24. Chen Q, Yan M, Cao Z, Li X, Zhang Y, Shi J, Feng G, Peng H, Zhang X, Zhang Y, et al (2016) Sperm tsRNAs contribute to intergenerational inheritance of an acquired metabolic disorder. Science 351: 397–400

25. Choi H, Wang Z & Dean J (2021) Sperm acrosome overgrowth and infertility in mice lacking chromosome 18 pachytene piRNA. PLOS Genet 17: e1009485

26. Conine CC, Sun F, Song L, Rivera-Pérez JA & Rando OJ (2018) Small RNAs gained during epididymal transit of sperm are essential for embryonic development in mice. Dev Cell 46: 470–480.e3

27. Conine CC, Sun F, Song L, Rivera-Pérez JA & Rando OJ (2020) Sperm Head Preparation and Genetic Background Affect Caput Sperm ICSI Embryo Viability: Cauda-Enriched miRNAs Only Essential in Specific Conditions. Dev Cell 55: 677–678

28. Crooks GE, Hon G, Chandonia J-M & Brenner SE (2004) WebLogo: a sequence logo generator. Genome Res 14: 1188–1190

29. Deng W & Lin H (2002) miwi, a Murine Homolog of piwi, Encodes a Cytoplasmic Protein Essential for Spermatogenesis. Dev Cell 2: 819–830

30. Diederichs S & Haber DA (2007) Dual role for argonautes in microRNA processing and posttranscriptional regulation of microRNA expression. Cell 131: 1097–1108

31. Dobin A, Davis CA, Schlesinger F, Drenkow J, Zaleski C, Jha S, Batut P, Chaisson M & Gingeras TR (2013) STAR: ultrafast universal RNA-seq aligner. Bioinformatics 29: 15–21

32. Esteves SC, Roque M, Bedoschi G, Haahr T & Humaidan P (2018) Intracytoplasmic sperm injection for male infertility and consequences for offspring. Nat Rev Urol 15: 535–562

33. Fabry MH, Falconio FA, Joud F, Lythgoe EK, Czech B & Hannon GJ (2021) Maternally inherited piRNAs direct transient heterochromatin formation at active transposons during early Drosophila embryogenesis. eLife 10: e68573

34. Foresta C, Rossato M, Mioni R & Zorzi M (1992) Progesterone induces capacitation in human spermatozoa. Andrologia 24: 33–35

35. Fu Y, Wu P-H, Beane T, Zamore PD & Weng Z (2018) Elimination of PCR duplicates in RNA-seq and small RNA-seq using unique molecular identifiers. BMC Genomics 19: 531

36. Gainetdinov I, Colpan C, Cecchini K, Arif A, Jouravleva K, Albosta P, Vega-Badillo J, Lee Y, Özata DM & Zamore PD (2021) Terminal modification, sequence, length, and PIWI-protein identity determine piRNA stability. Mol Cell 81: 4826–4842.e8

37. Gainetdinov I, Vega-Badillo J, Cecchini K, Bagci A, Colpan C, De D, Bailey S, Arif A, Wu P-H, MacRae IJ, et al (2023) Relaxed targeting rules help PIWI proteins silence transposons. Nature 619: 394–402

38. Gao H, Wen H, Cao C, Dong D, Yang C, Xie S, Zhang J, Huang X, Huang X, Yuan S, et al (2019) Overexpression of MicroRNA-10a in Germ Cells Causes Male Infertility by Targeting Rad51 in Mouse and Human. Front Physiol 10: 765

39. Georgiadis AP, Kishore A, Zorrilla M, Jaffe TM, Sanfilippo JS, Volk E, Rajkovic A & Yatsenko AN (2015) High quality RNA in semen and sperm: isolation, analysis and potential application in clinical testing. J Urol 193: 352–359

40. Gerstberger S, Hafner M & Tuschl T (2014) A census of human RNA-binding proteins. Nat Rev Genet 15: 829–845

41. Gervasi MG & Visconti PE (2017) Molecular changes and signaling events occurring in spermatozoa during epididymal maturation. Andrology 5: 204–218

42. Ghildiyal M & Zamore PD (2009) Small silencing RNAs: an expanding universe. Nat Rev Genet 10: 94–108

43. Girard A, Sachidanandam R, Hannon GJ & Carmell MA (2006) A germline-specific class of small RNAs binds mammalian Piwi proteins. Nature 442: 199–202

44. Gòdia M, Mayer FQ, Nafissi J, Castelló A, Rodríguez-Gil JE, Sánchez A & Clop A (2018) A technical assessment of the porcine ejaculated spermatozoa for a sperm-specific RNA-seq analysis. Syst Biol Reprod Med 64: 291–303

45. Goh WSS, Falciatori I, Tam OH, Burgess R, Meikar O, Kotaja N, Hammell M & Hannon GJ (2015) piRNA-directed cleavage of meiotic transcripts regulates spermatogenesis. Genes Dev 29: 1032–1044

46. Gong Y, Li L, Qian Y, Lu T, Zhang Z, Jiang L, Liu G, Cui M, Li S, Li Z, et al (2025) PIWIL3-piRNA Pathway Is Essential for Rabbit Oogenesis and Embryogenesis via Broad Regulation of the Transcriptome and Proteome. doi:10.1101/2025.10.23.684072 [PREPRINT]

47. Gong Y, Shi S, Li L, Qian Y, Lu T, Zhang Z, Jiang L, Liu G, Cui M, Li S, et al (2026) PIWIL3-piRNA pathway controls rabbit oogenesis and embryogenesis via broad regulation of the transcriptome and proteome. Nat Commun

48. González-González E, López-Casas PP & del Mazo J (2008) The expression patterns of genes involved in the RNAi pathways are tissue-dependent and differ in the germ and somatic cells of mouse testis. Biochim Biophys Acta 1779: 306–311

49. Gou L-T, Kang J-Y, Dai P, Wang X, Li F, Zhao S, Zhang M, Hua M-M, Lu Y, Zhu Y, et al (2017) Ubiquitination-Deficient Mutations in Human Piwi Cause Male Infertility by Impairing Histone-to-Protamine Exchange during Spermiogenesis. Cell 169: 1090–1104.e13

50. Grentzinger T, Oberlin S, Schott G, Handler D, Svozil J, Barragan-Borrero V, Humbert A, Duharcourt S, Brennecke J & Voinnet O (2020) A universal method for the rapid isolation of all known classes of functional silencing small RNAs. Nucleic Acids Res 48: e79

51. Gu H, Lian B, Yuan Y, Kong C, Li Y, Liu C & Qi Y (2022) A 5’ tRNA-Ala-derived small RNA regulates anti-fungal defense in plants. Sci China Life Sci 65: 1–15

52. Gustafsson HT, Ferguson L, Galan C, Yu T, Upton H, Kaymak E, Weng Z, Collins K & Rando OJ (2025) Deep sequencing of yeast and mouse tRNAs and tRNA fragments using OTTR. eLife 14: e77616

53. Hasuwa H, Iwasaki YW, Au Yeung WK, Ishino K, Masuda H, Sasaki H & Siomi H (2021) Production of functional oocytes requires maternally expressed PIWI genes and piRNAs in golden hamsters. Nat Cell Biol 23: 1002–1012

54. Hentze MW, Castello A, Schwarzl T & Preiss T (2018) A brave new world of RNA-binding proteins. Nat Rev Mol Cell Biol 19: 327–341

55. Hong J & Gresham D (2017) Incorporation of unique molecular identifiers in TruSeq adapters improves the accuracy of quantitative sequencing. BioTechniques 63: 221–226

56. Hutcheon K, McLaughlin EA, Stanger SJ, Bernstein IR, Dun MD, Eamens AL & Nixon B (2017) Analysis of the small non-protein-coding RNA profile of mouse spermatozoa reveals specific enrichment of piRNAs within mature spermatozoa. RNA Biol 14: 1776–1790

57. Isacson S, Karlsson K, Zalavary S, Asratian A, Kugelberg U, Liffner S & Öst A (2025) Small RNA in sperm–Paternal contributions to human embryo development. Nat Commun 16: 6571

58. Ishino K, Hasuwa H, Yoshimura J, Iwasaki YW, Nishihara H, Seki NM, Hirano T, Tsuchiya M, Ishizaki H, Masuda H, et al (2021) Hamster PIWI proteins bind to piRNAs with stage-specific size variations during oocyte maturation. Nucleic Acids Res 49: 2700–2720

59. Johnson GD, Mackie P, Jodar M, Moskovtsev S & Krawetz SA (2015) Chromatin and extracellular vesicle associated sperm RNAs. Nucleic Acids Res 43: 6847–6859

60. Johnson GD, Sendler E, Lalancette C, Hauser R, Diamond MP & Krawetz SA (2011) Cleavage of rRNA ensures translational cessation in sperm at fertilization. Mol Hum Reprod 17: 721–726

61. Käll L, Storey JD & Noble WS (2008) Non-parametric estimation of posterior error probabilities associated with peptides identified by tandem mass spectrometry. Bioinformatics 24: i42–48

62. Kivioja T, Vähärautio A, Karlsson K, Bonke M, Enge M, Linnarsson S & Taipale J (2011) Counting absolute numbers of molecules using unique molecular identifiers. Nat Methods 9: 72–74

63. Kojima K, Kuramochi-Miyagawa S, Chuma S, Tanaka T, Nakatsuji N, Kimura T & Nakano T (2009) Associations between PIWI proteins and TDRD1/MTR-1 are critical for integrated subcellular localization in murine male germ cells. Genes Cells Devoted Mol Cell Mech 14: 1155–1165

64. König L, Guggenberger V, Eleftheriou K, Pinter Z, Marotto A, Kreutz CR, Wossidlo M, Marchand V, Motorin Y & Schaefer MR (2025) Copy number determination of sperm-borne small RNAs implied in the intergenerational inheritance of metabolic syndromes. RNA 31: 1041–1052

65. Korneev D, Merriner DJ, Gervinskas G, de Marco A & O’Bryan MK (2021) New Insights Into Sperm Ultrastructure Through Enhanced Scanning Electron Microscopy. Front Cell Dev Biol 9: 672592

66. Kotaja N & Sassone-Corsi P (2007) The chromatoid body: a germ-cell-specific RNA-processing centre. Nat Rev Mol Cell Biol 8: 85–90

67. Kozomara A, Birgaoanu M & Griffiths-Jones S (2019) miRBase: from microRNA sequences to function. Nucleic Acids Res 47: D155–D162

68. Krawetz SA, Kruger A, Lalancette C, Tagett R, Anton E, Draghici S & Diamond MP (2011) A survey of small RNAs in human sperm. Hum Reprod 26: 3401–3412

69. de Kretser DM, Loveland KL, Meinhardt A, Simorangkir D & Wreford N (1998) Spermatogenesis. Hum Reprod 13 Suppl 1: 1–8

70. Kugler J-M, Chen Y-W, Weng R & Cohen SM (2013) Maternal Loss of miRNAs Leads to Increased Variance in Primordial Germ Cell Numbers in *Drosophila melanogaster*. G3 Genes Genomes Genet 3: 1573–1576

71. Kuramochi-Miyagawa S, Watanabe T, Gotoh K, Totoki Y, Toyoda A, Ikawa M, Asada N, Kojima K, Yamaguchi Y, Ijiri TW, et al (2008) DNA methylation of retrotransposon genes is regulated by Piwi family members MILI and MIWI2 in murine fetal testes. Genes Dev 22: 908–917

72. Langmead B, Trapnell C, Pop M & Salzberg SL (2009) Ultrafast and memory-efficient alignment of short DNA sequences to the human genome. Genome Biol 10: R25

73. Lei Y, Zhang X, Xu Q, Liu S, Li C, Jiang H, Lin H, Kong E, Liu J, Qi S, et al (2021) Autophagic elimination of ribosomes during spermiogenesis provides energy for flagellar motility. Dev Cell 56: 2313–2328.e7

74. Li XZ, Roy CK, Dong X, Bolcun-Filas E, Wang J, Han BW, Xu J, Moore MJ, Schimenti JC, Weng Z, et al (2013) An ancient transcription factor initiates the burst of piRNA production during early meiosis in mouse testes. Mol Cell 50: 67–81

75. Li Y, Li R-H, Ran M-X, Zhang Y, Liang K, Ren Y-N, He W-C, Zhang M, Zhou G-B, Qazi IH, et al (2018) High throughput small RNA and transcriptome sequencing reveal capacitation-related microRNAs and mRNA in boar sperm. BMC Genomics 19

76. Li Z, Liu X, Zhang Y, Li Y, Zhou L & Yuan S (2024) FBXO24 modulates mRNA alternative splicing and MIWI degradation and is required for normal sperm formation and male fertility. eLife 12: RP91666

77. Liao J-Y, Yang B, Shi C-P, Deng W-X, Deng J-S, Cen M-F, Zheng B-Q, Zhan Z-L, Liang Q-L, Wang J-E, et al (2025) RBPWorld for exploring functions and disease associations of RNA-binding proteins across species. Nucleic Acids Res 53: D220–D232

78. Lim SL, Qu ZP, Kortschak RD, Lawrence DM, Geoghegan J, Hempfling A-L, Bergmann M, Goodnow CC, Ormandy CJ, Wong L, et al (2015) HENMT1 and piRNA Stability Are Required for Adult Male Germ Cell Transposon Repression and to Define the Spermatogenic Program in the Mouse. PLoS Genet 11: e1005620

79. Liu T, Huang Y, Liu J, Zhao Y, Jiang L, Huang Q, Cheng W & Guo L (2013) MicroRNA-122 influences the development of sperm abnormalities from human induced pluripotent stem cells by regulating TNP2 expression. Stem Cells Dev 22: 1839–1850

80. Liu W-M, Pang RTK, Chiu PCN, Wong BPC, Lao K, Lee K-F & Yeung WSB (2012) Sperm-borne microRNA-34c is required for the first cleavage division in mouse. Proc Natl Acad Sci 109: 490–494

81. Loubalova Z, Fulka H, Horvat F, Pasulka J, Malik R, Hirose M, Ogura A & Svoboda P (2021) Formation of spermatogonia and fertile oocytes in golden hamsters requires piRNAs. Nat Cell Biol 23: 992–1001

82. Love MI, Huber W & Anders S (2014) Moderated estimation of fold change and dispersion for RNA-seq data with DESeq2. Genome Biol 15: 550

83. Lv X, Xiao W, Lai Y, Zhang Z, Zhang H, Qiu C, Hou L, Chen Q, Wang D, Gao Y, et al (2023) The non-redundant functions of PIWI family proteins in gametogenesis in golden hamsters. Nat Commun 14: 5267

84. Martin M (2011) Cutadapt removes adapter sequences from high-throughput sequencing reads. EMBnet.journal 17: 10

85. Matsumoto K & Wolffe AP (1998) Gene regulation by Y-box proteins: coupling control of transcription and translation. Trends Cell Biol 8: 318–323

86. Meikar O, Vagin VV, Chalmel F, Sõstar K, Lardenois A, Hammell M, Jin Y, Da Ros M, Wasik KA, Toppari J, et al (2014) An atlas of chromatoid body components. RNA 20: 483–495

87. Modzelewski AJ, Holmes RJ, Hilz S, Grimson A & Cohen PE (2012) AGO4 regulates entry into meiosis and influences silencing of sex chromosomes in the male mouse germline. Dev Cell 23: 251–264

88. Mortimer D (2018) The functional anatomy of the human spermatozoon: relating ultrastructure and function. Mol Hum Reprod 24: 567–592

89. Motorin Y & Marchand V (2018) Detection and Analysis of RNA Ribose 2’-O-Methylations: Challenges and Solutions. Genes 9: 642

90. Nesvizhskii AI, Keller A, Kolker E & Aebersold R (2003) A statistical model for identifying proteins by tandem mass spectrometry. Anal Chem 75: 4646–4658

91. Nixon B, Stanger SJ, Mihalas BP, Reilly JN, Anderson AL, Tyagi S, Holt JE & McLaughlin EA (2015) The microRNA signature of mouse spermatozoa is substantially modified during epididymal maturation. Biol Reprod 93: 91

92. O’Donnell L, Nicholls PK, O’Bryan MK, McLachlan RI & Stanton PG (2011) Spermiation: The process of sperm release. Spermatogenesis 1: 14–35

93. Ord J, Heath PR, Fazeli A & Watt PJ (2020) Paternal effects in a wild-type zebrafish implicate a role of sperm-derived small RNAs. Mol Ecol 29: 2722–2735

94. Ozata DM, Gainetdinov I, Zoch A, O’Carroll D & Zamore PD (2019) PIWI-interacting RNAs: small RNAs with big functions. Nat Rev Genet 20: 89–108

95. Padhiar NH, Katneni U, Komar AA, Motorin Y & Kimchi-Sarfaty C (2024) Advances in methods for tRNA sequencing and quantification. Trends Genet TIG 40: 276–290

96. Pantano L, Jodar M, Bak M, Ballescà JL, Tommerup N, Oliva R & Vavouri T (2015) The small RNA content of human sperm reveals pseudogene-derived piRNAs complementary to protein-coding genes. RNA 21: 1085–1095

97. Parisien M, Wang X & Pan T (2013) Diversity of human tRNA genes from the 1000-genomes project. RNA Biol 10: 1853–1867

98. Pearson BL, Posselt LP, Henzel KS & Ehninger D (2018) Limited efficacy of somatic cell lysis buffer to purify laboratory mouse sperm. Epigenomics 10: 689–694

99. Peng H, Shi J, Zhang Y, Zhang H, Liao S, Li W, Lei L, Han C, Ning L, Cao Y, et al (2012) A novel class of tRNA-derived small RNAs extremely enriched in mature mouse sperm. Cell Res 22: 1609–1612

100. Piriyapongsa J, Mariño-Ramírez L & Jordan IK (2007) Origin and evolution of human microRNAs from transposable elements. Genetics 176: 1323–1337

101. Prathalingam NS, Holt WW, Revell SG, Jones S & Watson PF (2006) The precision and accuracy of six different methods to determine sperm concentration. J Androl 27: 257–262

102. Puga Molina LC, Luque GM, Balestrini PA, Marín-Briggiler CI, Romarowski A & Buffone MG (2018) Molecular basis of human sperm capacitation. Front Cell Dev Biol 6: 72

103. Quarato P, Singh M, Cornes E, Li B, Bourdon L, Mueller F, Didier C & Cecere G (2021) Germline inherited small RNAs facilitate the clearance of untranslated maternal mRNAs in C. elegans embryos. Nat Commun 12: 1441

104. Quinn P, Kerin JF & Warnes GM (1985) Improved pregnancy rate in human in vitro fertilization with the use of a medium based on the composition of human tubal fluid. Fertil Steril 44: 493–498

105. Raad N, Fernandez-Rodriguez C, Pandey RR, Mohammed I, Uchikawa E, Burger F, Homolka D & Pillai RS (2026) Structure of the MIWI endoribonuclease bound to pachytene piRNAs from mouse testes. Cell Rep 45: 116804

106. Rando OJ (2012) Daddy issues: paternal effects on phenotype. Cell 151: 702–708

107. Rassoulzadegan M, Grandjean V, Gounon P, Vincent S, Gillot I & Cuzin F (2006) RNA-mediated non-mendelian inheritance of an epigenetic change in the mouse. Nature 441: 469–474

108. Rengan AK, Agarwal A, van der Linde M & du Plessis SS (2012) An investigation of excess residual cytoplasm in human spermatozoa and its distinction from the cytoplasmic droplet. Reprod Biol Endocrinol RBE 10: 92

109. Reuter M, Berninger P, Chuma S, Shah H, Hosokawa M, Funaya C, Antony C, Sachidanandam R & Pillai RS (2011) Miwi catalysis is required for piRNA amplification-independent LINE1 transposon silencing. Nature 480: 264–267

110. Rieger I, Weintraub G, Lev I, Goldstein K, Bar-Zvi D, Anava S, Gingold H, Shaham S & Rechavi O (2023) Nucleus-independent transgenerational small RNA inheritance in Caenorhabditis elegans. Sci Adv 9: eadj8618

111. Rinaldi VD, Donnard E, Gellatly K, Rasmussen M, Kucukural A, Yukselen O, Garber M, Sharma U & Rando OJ (2020) An atlas of cell types in the mouse epididymis and vas deferens. eLife 9: e55474

112. Rodgers AB, Morgan CP, Leu NA & Bale TL (2015) Transgenerational epigenetic programming via sperm microRNA recapitulates effects of paternal stress. Proc Natl Acad Sci 112: 13699–13704

113. Roovers EF, Rosenkranz D, Mahdipour M, Han C-T, He N, Chuva de Sousa Lopes SM, van der Westerlaken LAJ, Zischler H, Butter F, Roelen BAJ, et al (2015) Piwi proteins and piRNAs in mammalian oocytes and early embryos. Cell Rep 10: 2069–2082

114. Rouget C, Papin C, Boureux A, Meunier A-C, Franco B, Robine N, Lai EC, Pelisson A & Simonelig M (2010) Maternal mRNA deadenylation and decay by the piRNA pathway in the early Drosophila embryo. Nature 467: 1128–1132

115. Schreier J, Dietz S, Boermel M, Oorschot V, Seistrup A-S, de Jesus Domingues AM, Bronkhorst AW, Nguyen DAH, Phillis S, Gleason EJ, et al (2022) Membrane-associated cytoplasmic granules carrying the Argonaute protein WAGO-3 enable paternal epigenetic inheritance in Caenorhabditis elegans. Nat Cell Biol 24: 217–229

116. Schuster A, Tang C, Xie Y, Ortogero N, Yuan S & Yan W (2016) SpermBase: A Database for Sperm-Borne RNA Contents. Biol Reprod 95: 99–99

117. Sellem E, Marthey S, Rau A, Jouneau L, Bonnet A, Perrier J-P, Fritz S, Le Danvic C, Boussaha M, Kiefer H, et al (2020) A comprehensive overview of bull sperm-borne small non-coding RNAs and their diversity across breeds. Epigenetics Chromatin 13: 19

118. Sharma U, Conine CC, Shea JM, Boskovic A, Derr AG, Bing XY, Belleannee C, Kucukural A, Serra RW, Sun F, et al (2016) Biogenesis and function of tRNA fragments during sperm maturation and fertilization in mammals. Science 351: 391–396

119. Sharma U, Sun F, Conine CC, Reichholf B, Kukreja S, Herzog VA, Ameres SL & Rando OJ (2018) Small RNAs Are Trafficked from the Epididymis to Developing Mammalian Sperm. Dev Cell 46: 481–494.e6

120. Shi J, Fok KL, Dai P, Qiao F, Zhang M, Liu H, Sang M, Ye M, Liu Y, Zhou Y, et al (2021a) Spatio-temporal landscape of mouse epididymal cells and specific mitochondria-rich segments defined by large-scale single-cell RNA-seq. Cell Discov 7: 34

121. Shi J, Zhang Y, Tan D, Zhang X, Yan M, Zhang Y, Franklin R, Shahbazi M, Mackinlay K, Liu S, et al (2021b) PANDORA-seq expands the repertoire of regulatory small RNAs by overcoming RNA modifications. Nat Cell Biol 23: 424–436

122. Short AK, Yeshurun S, Powell R, Perreau VM, Fox A, Kim JH, Pang TY & Hannan AJ (2017) Exercise alters mouse sperm small noncoding RNAs and induces a transgenerational modification of male offspring conditioned fear and anxiety. Transl Psychiatry 7: e1114

123. Skerrett-Byrne DA, Anderson AL, Bromfield EG, Bernstein IR, Mulhall JE, Schjenken JE, Dun MD, Humphrey SJ & Nixon B (2022) Global profiling of the proteomic changes associated with the post-testicular maturation of mouse spermatozoa. Cell Rep 41: 111655

124. Smith T, Heger A & Sudbery I (2017) UMI-tools: modeling sequencing errors in Unique Molecular Identifiers to improve quantification accuracy. Genome Res 27: 491–499

125. Song R, Walentek P, Sponer N, Klimke A, Lee JS, Dixon G, Harland R, Wan Y, Lishko P, Lize M, et al (2014) miR-34/449 miRNAs are required for motile ciliogenesis by repressing cp110. Nature 510: 115–120

126. Sprando RL & Russell LD (1987) Comparative study of cytoplasmic elimination in spermatids of selected mammalian species. Am J Anat 178: 72–80

127. Stanger SJ, Bernstein IR, Anderson AL, Hutcheon K, Dun MD, Eamens AL & Nixon B (2020) The abundance of a transfer RNA-derived RNA fragment small RNA subpopulation is enriched in cauda spermatozoa. ExRNA 2: 17

128. Stoeckius M, Grün D & Rajewsky N (2014) Paternal RNA contributions in the Caenorhabditis elegans zygote. EMBO J 33: 1740–1750

129. Suh N, Baehner L, Moltzahn F, Melton C, Shenoy A, Chen J & Blelloch R (2010) MicroRNA function is globally suppressed in mouse oocytes and early embryos. Curr Biol CB 20: 271–277

130. Sun J, Philpott M, Loi D, Li S, Monteagudo-Mesas P, Hoffman G, Robson J, Mehta N, Gamble V, Brown T, et al (2024) Correcting PCR amplification errors in unique molecular identifiers to generate accurate numbers of sequencing molecules. Nat Methods 21: 401–405

131. Sun YH, Wang A, Song C, Shankar G, Srivastava RK, Au KF & Li XZ (2021) Single-molecule long-read sequencing reveals a conserved intact long RNA profile in sperm. Nat Commun 12: 1361

132. Szklarczyk D, Nastou K, Koutrouli M, Kirsch R, Mehryary F, Hachilif R, Hu D, Peluso ME, Huang Q, Fang T, et al (2025) The STRING database in 2025: protein networks with directionality of regulation. Nucleic Acids Res 53: D730–D737

133. Sztein JM, Farley JS & Mobraaten LE (2000) In vitro fertilization with cryopreserved inbred mouse sperm. Biol Reprod 63: 1774–1780

134. Takemoto N, Yoshimura T, Miyazaki S, Tashiro F & Miyazaki J (2016) Gtsf1l and Gtsf2 Are Specifically Expressed in Gonocytes and Spermatids but Are Not Essential for Spermatogenesis. PloS One 11: e0150390

135. Tan M, van Tol HTA, Mokry M, Stout TAE & Roelen BAJ (2020) Microinjection induces changes in the transcriptome of bovine oocytes. Sci Rep 10: 11211

136. Tang F, Kaneda M, O’Carroll D, Hajkova P, Barton SC, Sun YA, Lee C, Tarakhovsky A, Lao K & Surani MA (2007) Maternal microRNAs are essential for mouse zygotic development. Genes Dev 21: 644–648

137. The UniProt Consortium, Bateman A, Martin M-J, Orchard S, Magrane M, Adesina A, Ahmad S, Bowler-Barnett EH, Bye-A-Jee H, Carpentier D, et al (2025) UniProt: the Universal Protein Knowledgebase in 2025. Nucleic Acids Res 53: D609–D617

138. Tomar A, Gomez-Velazquez M, Gerlini R, Comas-Armangué G, Makharadze L, Kolbe T, Boersma A, Dahlhoff M, Burgstaller JP, Lassi M, et al (2024) Epigenetic inheritance of diet-induced and sperm-borne mitochondrial RNAs. Nature 630: 720–727

139. Tomlinson M, Turner J, Powell G & Sakkas D (2001) One-step disposable chambers for sperm concentration and motility assessment: how do they compare with the World Health Organization’s recommended methods? Hum Reprod 16: 121–124

140. Tuttle AH, Philip VM, Chesler EJ & Mogil JS (2018) Comparing phenotypic variation between inbred and outbred mice. Nat Methods 15: 994–996

141. Tyanova S, Temu T & Cox J (2016) The MaxQuant computational platform for mass spectrometry-based shotgun proteomics. Nat Protoc 11: 2301–2319

142. Vagin VV, Wohlschlegel J, Qu J, Jonsson Z, Huang X, Chuma S, Girard A, Sachidanandam R, Hannon GJ & Aravin AA (2009) Proteomic analysis of murine Piwi proteins reveals a role for arginine methylation in specifying interaction with Tudor family members. Genes Dev 23: 1749–1762

143. Visconti PE, Bailey JL, Moore GD, Pan D, Olds-Clarke P & Kopf GS (1995a) Capacitation of mouse spermatozoa. I. Correlation between the capacitation state and protein tyrosine phosphorylation. Development 121: 1129–1137

144. Visconti PE, Galantino-Homer H, Moore GD, Bailey JL, Ning X, Fornes M & Kopf GS (1998) The molecular basis of sperm capacitation. J Androl 19: 242–248

145. Visconti PE, Moore GD, Bailey JL, Leclerc P, Connors SA, Pan D, Olds-Clarke P & Kopf GS (1995b) Capacitation of mouse spermatozoa. II. Protein tyrosine phosphorylation and capacitation are regulated by a cAMP-dependent pathway. Development 121: 1139–1150

146. Walker WH (2022) Regulation of mammalian spermatogenesis by miRNAs. Semin Cell Dev Biol 121: 24–31

147. Wang H, Wang Z, Zhou T, Morris D, Chen S, Li M, Wang Y, Zheng H, Fu W & Yan W (2023) Small RNA shuffling between murine sperm and their cytoplasmic droplets during epididymal maturation. Dev Cell 58: 779–790.e4

148. Wang M, Du Y, Gao S, Wang Z, Qu P, Gao Y, Wang J, Liu Z, Zhang J, Zhang Y, et al (2021a) Sperm-borne miR-202 targets SEPT7 and regulates first cleavage of bovine embryos via cytoskeletal remodeling. Development 148: dev189670

149. Wang Y, Chen Z-P, Hu H, Lei J, Zhou Z, Yao B, Chen L, Liang G, Zhan S, Zhu X, et al (2021b) Sperm microRNAs confer depression susceptibility to offspring. Sci Adv 7: eabd7605

150. Wang Y, Yamauchi Y, Wang Z, Zheng H, Yanagimachi R, Ward MA & Yan W (2020) Both Cauda and Caput Epididymal Sperm Are Capable of Supporting Full-Term Development in FVB and CD-1 Mice. Dev Cell 55: 675–676

151. Winter J & Diederichs S (2011) Argonaute proteins regulate microRNA stability: Increased microRNA abundance by Argonaute proteins is due to microRNA stabilization. RNA Biol 8: 1149–1157

152. Wu P-H, Fu Y, Cecchini K, Özata DM, Arif A, Yu T, Colpan C, Gainetdinov I, Weng Z & Zamore PD (2020) The evolutionarily conserved piRNA-producing locus pi6 is required for male mouse fertility. Nat Genet 52: 728–739

153. Yang M, Oatley MJ, Miao D, Maddison LA & Oatley JM (2026) Biogenesis and migration of the sperm cytoplasmic droplet require ARRDC5 and TEX38. Cell Rep 45: 116731

154. Yu T, Fan K, Özata DM, Zhang G, Fu Y, Theurkauf WE, Zamore PD & Weng Z (2021) Long first exons and epigenetic marks distinguish conserved pachytene piRNA clusters from other mammalian genes. Nat Commun 12: 73

155. Yuan S, Liu Y, Peng H, Tang C, Hennig GW, Wang Z, Wang L, Yu T, Klukovich R, Zhang Y, et al (2019) Motile cilia of the male reproductive system require miR-34/miR-449 for development and function to generate luminal turbulence. Proc Natl Acad Sci 116: 3584–3593

156. Yuan S, Tang C, Zhang Y, Wu J, Bao J, Zheng H, Xu C & Yan W (2015) mir-34b/c and mir-449a/b/c are required for spermatogenesis, but not for the first cleavage division in mice. Biol Open 4: 212–223

157. Zhang H, Zhang F, Chen Q, Li M, Lv X, Xiao Y, Zhang Z, Hou L, Lai Y, Zhang Y, et al (2021) The piRNA pathway is essential for generating functional oocytes in golden hamsters. Nat Cell Biol 23: 1013–1022

158. Zhao S, Gou L-T, Zhang M, Zu L-D, Hua M-M, Hua Y, Shi H-J, Li Y, Li J, Li D, et al (2013) piRNA-triggered MIWI ubiquitination and removal by APC/C in late spermatogenesis. Dev Cell 24: 13–25

159. Zheng H, Stratton CJ, Morozumi K, Jin J, Yanagimachi R & Yan W (2007) Lack of Spem1 causes aberrant cytoplasm removal, sperm deformation, and male infertility. Proc Natl Acad Sci U S A 104: 6852–6857

160. Zhu T, Liao K, Zhou R, Xia C & Xie W (2020) ATAC-seq with unique molecular identifiers improves quantification and footprinting. Commun Biol 3: 675

